# Comprehensive analysis of kill switch toxins in plant-beneficial *Pseudomonas fluorescens* reveals drivers of lethality, stability, and escape

**DOI:** 10.1101/2022.07.18.500305

**Authors:** Tiffany M. Halvorsen, Dante P. Ricci, Dan M. Park, Yongqin Jiao, Mimi C. Yung

## Abstract

Kill switches provide a biocontainment strategy in which unwanted growth of an engineered microorganism is prevented by expression of a toxin gene. A major challenge in kill switch engineering is balancing evolutionary stability with robust cell killing activity in application relevant host strains. Understanding host-specific containment dynamics and modes of failure helps to develop potent yet stable kill switches. To guide the design of robust kill switches in the agriculturally relevant strain *Pseudomonas fluorescens* SBW25, we present a comparison of lethality, stability, and genetic escape of eight different toxic effectors in the presence of their cognate inactivators (i.e., toxin-antitoxin modules, polymorphic exotoxin-immunity systems, restriction endonuclease-methyltransferase pair). We find that cell killing capacity and evolutionary stability are inversely correlated and dependent on the level of protection provided by the inactivator gene. Decreasing the proteolytic stability of the inactivator protein can increase cell killing capacity, but at the cost of long-term circuit stability. By comparing toxins within the same genetic context, we determine that modes of genetic escape increase with circuit complexity and are driven by toxin activity, the protective capacity of the inactivator, and the presence of mutation-prone sequences within the circuit. Collectively, our study reveals that circuit complexity, toxin choice, inactivator stability, and DNA sequence design are powerful drivers of kill switch stability and valuable targets for optimization of biocontainment systems.

## Introduction

Synthetic biology promises to sustainably produce commodity chemicals^1–3^, treat human disease^4,5^, steward the environment^6,7^, and sustain agriculture^8^. Each of these tasks exploits genetically engineered microorganisms (GEMs), but the unintended release of GEMs or their recombinant DNA could disturb native ecosystems in unpredictable ways. Biocontainment strategies are therefore needed to limit unforeseen environmental disturbance. “Kill switch” circuits are a biocontainment method that prohibit GEM growth by regulated induction of a toxin gene via an environmentally responsive transcription factor^9,10^. The genetic components of kill switches are inherently modular, meaning that a wealth of options exist for regulators and toxins. Kill switches that respond to pH^11^, temperature^12,13^, and CO2^14^ have been devised, and the additional innate modularity of transcription factors means DNA-binding and small molecule-sensing domains can be shuffled to enable circuit customization^10^. Many well-characterized protein toxins also exist for driving containment, including toxin-antitoxin (TA) systems, proteases, polymorphic exotoxins, anti-phage restriction or CRISPR-associated endonucleases, and phage lysis genes^9,10,13,15–19^. To date, however, the vast majority of kill switches have been developed in *Escherichia coli*^10,11,13,18–22^, and tested under ideal laboratory conditions (i.e., in rich media). While kill switch designs have been developed in *Pseudomonas putida* and *Saccharomyces cerevisiae*^9,16,17,20,23–25^, these studies reveal that porting of kill switch parts from *E. coli* into alternative chassis strains is challenging, resulting in unpredictable changes in expression and fitness^9,16^. Moreover, GEMs developed for environmental applications will ultimately be deployed into uncontrolled environments, causing changes to growth rate and metabolic regulation that could affect circuit function and failure ^26^. Therefore, it is necessary to develop and study kill switches and their component parts (e.g., toxins, regulators) directly in application-specific GEM chassis and to challenge their robustness under different growth conditions.

When porting kill switches into new strains, a major challenge is striking a balance between sufficient toxicity in the non-permissive state (“on”) with near wild-type fitness in the permissive state (“off”). Promoter leakiness even in the presence of a transcriptional repressor typically causes decreased fitness in the off-state and could select for inactivating mutations that render the GEM uncontained. Recently, Silver and colleagues devised a method to reduce this fitness defect by including an antitoxin (*ccdA*) into a circuit designed with a Type II toxin (*ccdB*)^13^. In addition to reducing fitness cost, including the antitoxin to drive the circuit’s off-state also creates an opportunity to induce cell death by removal of a permissive signal rather than addition of a non-permissive signal, an ideal strategy for self-regulating environmental kill switches. However, this optimization strategy has yet to be explored in other chassis outside of *E. coli* or with other toxin systems other than Type II TA systems. It is unclear how much of a barrier antitoxins or other toxin inactivators (i.e., immunity proteins, methylases, small RNAs) would present when ported into new hosts. Therefore, a deeper investigation of the killing dynamics and optimization targets of kill switches in relevant hosts would help to guide the design of application-specific containment circuits that exert minimal fitness costs on the host strain.

Another major challenge associated with kill switch deployment in GEMs is escape from containment via mutational inactivation of any circuit component imparting a fitness cost on the host (e.g., promoter, toxin, regulator)^10,15,18^. Different host-specific drivers of genetic escape would alter the spectrum of mutations and specific sequences that lead to circuit inactivation (e.g., mobile element insertions; recombination-mediated deletions; replication errors) ^15,16,18,27,28^. Therefore, profiling escape mutants in specific chassis is important to understand these drivers of inactivation and ultimately identify strategies to prevent inactivation to achieve sufficiently long evolutionary stability^29^. Previous efforts to minimize genetic escape have opted to include redundant or multiple toxins in the design, which reduces the probability of total circuit inactivation^15,18^. An alternative strategy to reduce mutational escape is to eliminate routes by which the host strain accrues inactivating mutations, such as stress-response genes and RecA^10,16,18^. Finally, elimination of mobile elements from the host genome can greatly reduce escape frequency in some hosts^27,30^. Since the effectiveness of circuit stabilization strategies is host dependent, determining the major modes of escape in each host is critical for the development of stable kill switches.

Here, we set out to obtain a better understanding of host-specific drivers of kill switch functionality and stability in an environmental soil isolate, *Pseudomonas fluorescens* strain SBW25. Given the distinct lack of containment systems for plant growth promoting *P. fluorescens* and their propensity for use as GEMs in agriculture^31–34^, there is a need to develop biocontainment strategies for these strains. Toward this goal, we assessed a variety of toxin systems in *P. fluorescens*, including several Type II TA systems, a colicin endonuclease, a cytosolic toxin from a type VI secretion system, a restriction endonuclease, and a pore-forming Type I TA system. We tested each toxin in the presence of its cognate antitoxin or immunity gene (here, termed ‘inactivators’) to minimize fitness costs associated with leaky toxin expression. In doing so, we find that inactivator proteins are powerful determinants of kill switch functionality and stability. Finally, by analyzing modes and frequencies of escape mutations for each system, we unearth key drivers of genetic escape and important considerations for host-specific circuit design in *P. fluorescens*.

## Results & Discussion

### Established toxin-inactivator pairs are functional in *P. fluorescens*

To assess cell killing by toxin-inactivator systems from within the same genetic context in *P. fluorescens* SBW25, we chose to regulate toxin expression with the tightly controlled, cumate-inducible P_*cym*_-CymR promoter-repressor pair from *P. putida*, and inactivator expression with the leaky, isopropyl thiogalactoside (IPTG)-inducible P_*tac*_-LacI promoter-repressor pair to allow for relatively high basal expression to sequester leaked toxin^35^ (**Fig. 1A, Fig. S1, Table S1**). As *P. fluorescens* is a rhizobacterium, we measured cell killing in both rich (Luria-Bertani (LB)) and minimal M9 media with carbon sources relevant for growth in the rhizosphere (glucose, succinate, proline)^36,37^. As the major metric of toxicity throughout this study, we measured the fraction of total cells able to survive non-permissive induction and refer to this fraction as the survival ratio (i.e., colony forming units (CFU)/mL on non-permissive medium divided by CFU/mL on permissive medium). Importantly, colonies counted on non-permissive media may be a combination of genetically inactivated mutants (i.e., escapees) and cumate insensitive cells surviving induction due to insufficient toxicity.

**Figure 1.**
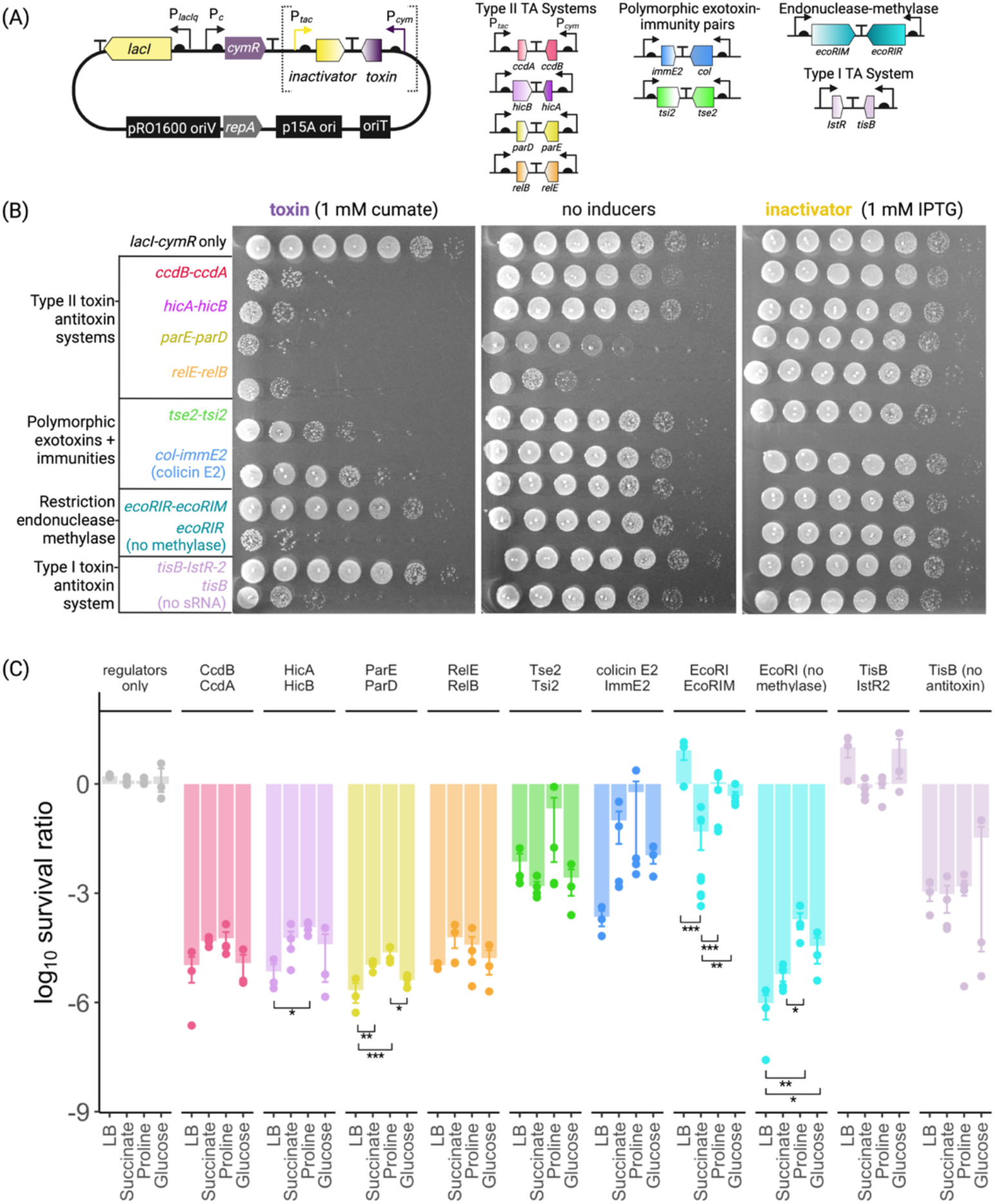
Kill switch efficacy for various toxins and their cognate inactivators under dual regulation in *P. fluorescens* SBW25. (A) Dual circuit plasmid used to screen toxin-inactivator pairs. Addition of IPTG induces inactivator expression by relieving P_*tac*_-bound LacI, while addition of cumate allows expression of the toxin via de-repression of P_*cym*_ by CymR. (B) Representative functional screen of all eight toxin-inactivator pairs in SBW25 tested in LB. Tenfold serial dilutions of SBW25 cultures, pre-grown with IPTG, were plated on LB-agar supplemented with 1 mM cumate (non-permissive, left), no inducers (to assess basal toxicity, middle), or 1 mM IPTG (permissive, right). EcoRI and TisB were also tested without their cognate inactivators as neither is toxic in the presence of its cognate inactivator. (C) Survival ratios of SBW25 carrying each kill switch on rich LB medium or minimal M9 medium with 20 mM of the indicated carbon source. Survival ratio is the CFU/mL on cumate compared to the total CFU/mL in the culture (counted on IPTG). Data represent calculations from three separate replicates ±SEM. One-way ANOVA tests were performed on each kill switch as a function of medium composition. Asterisks represent the following p-value cutoffs obtained from a post-hoc Tukey test: p>0.05 (none), p<0.05 (one), p<0.01 (two), p<0.001 (three).

The first panel of toxin-inactivator pairs that we tested under dual regulation include four Type II TA systems: two gyrase inhibitor systems (*ccdB-ccdA*, *parD-parE*) and two endoribonuclease systems (*hicA-hicB*, *relE-relB*). Non-permissive induction of all four TA systems results in low survival ratios (at least 10^-4^) regardless of growth conditions, indicating robust cell killing by each toxin under 1 mM cumate induction (**Fig. 1**). However, there is a significant increase in survival of SBW25 expressing *hicA-hicB* and *parE-parD* in minimal-proline compared to rich medium (**Fig. 1C**). This result highlights the importance of challenging kill switches in alternative growth conditions, where achieving optimal killing efficiency could require additional tuning of expression levels. We also find that fitness costs without induction vary among TA systems in our dual regulatory system: basal (uninduced) expression of *parE-parD* and *relE-relB* causes a greater fitness defect than that of *ccdB-ccdA* and *hicA-hicB* (**Fig. 1B**, middle panel). The high cost of basal *parE* and *relE* expression can be relieved by the addition of 1 mM IPTG, indicating that toxicity is specifically due to toxin expression and antitoxin over-expression is a viable means for reducing the fitness costs from leaked toxin (**Fig. 1B**). Moreover, removing IPTG from the inoculum culture leads to increased survival ratios for *parE* and *relE* (**Fig. S2**), suggesting that colonies surviving non-permissive selection by highly lethal circuits are genetically inactivated mutants that arise within the inoculum culture. We confirmed this result using sequencing, which we describe in detail in a subsequent mutational escape section.

Next, we examined highly lethal, bactericidal polymorphic exotoxins in our dual regulatory circuit (i.e., colicins, cargo of Type VI secretion systems). We opted to test a colicin DNase (E2) and an enzymatic Type VI toxin (Tse2) as they are cytosolic-acting and thus likely to be functional when internally expressed from a plasmid^38,39^. Moreover, colicin E2 could reduce the spread of recombinant DNA release following cell death. Although colicins are made by *E. coli* as secreted toxins with functional domains for translocating into target bacteria, the cytotoxic portion can be expressed alone to produce an intracellular toxin that maintains immunity binding propensity^40^. We were surprised to observe higher survival ratios compared to Type II TA systems upon expression of either *colE2* or *tse2* (10^-2^ – 10^-4^ with no significant difference across media conditions, **Fig. 1**). These toxins cannot be cloned without their respective immunity genes (*immE2*, *tsi2*), however, which indicates that they are highly toxic to SBW25 in the absence of their immunity proteins. Notably, antitoxins are targeted for rapid turnover by cellular proteases to allow free toxin to inhibit growth^41,42^. In contrast, immunity proteins have evolved to prevent autointoxication by irreversibly binding to their cognate toxin^38,43^. Therefore, immunity protection may present a greater barrier to cell killing than antitoxin protection. Indeed, over-expression of immunity proteins in the inoculum prior to non-permissive induction increases the survival ratio of SBW25 expressing either *tse2* or *colE2* (**Fig. S2**).

Finally, we tested a Type I pore-forming TA system (*tisB-IstR*) and an endonuclease-methyltransferase system (*ecoRIR-ecoRIM*) as kill switches. Interestingly, we find that both EcoRI and TisB can inhibit growth of SBW25 in LB, but only in the absence of the genes encoding their cognate inactivators (**Fig. 1B** last 4 rows). These results suggest that mRNA silencing and DNA methylation are more effective than protein antitoxins at preventing toxicity of their cognate toxins in our dual-regulatory expression system. Surprisingly, despite no evidence for toxicity in rich medium, we find that the *ecoRIR-ecoRIM* circuit inhibits growth of SBW25 when induced by cumate in minimal-succinate medium (~10^-3^ survival ratio) (**Fig. 1C**). We verified that this decreased survival is due to circuit toxicity by sequencing colonies from three separate lineages that survived non-permissive induction on both minimal-succinate and LB agar. Indeed, whereas three colonies isolated from LB-agar all returned intact circuits, those that were isolated from minimal-succinate contained large, circuit-inactivating deletions, indicating that they are escape mutants caused by an increase in the fitness cost of *ecoRIR* induction. We anticipate that a change in circuit output, growth rate, metabolic regulation, or a combination of these increases *ecoRIR-ecoRIM* circuit toxicity in minimal-succinate. Notably, in the absence of its antitoxin, TisB is less toxic to *P. fluorescens* than any of the Type II TA systems that we tested, indicated by 10-100 fold higher survival ratios for TisB that do not significantly change when tested in minimal media (**Fig. 1B-C**). In contrast, EcoRI in the absence of a methyltransferase is as toxic as the dual-regulated TA systems. EcoRI toxicity also decreases in minimal media, regardless of carbon source, which is consistent with the decreased toxicity of *hicA-hicB* and *parE-parD* in minimal media. Therefore, some circuits with low survival ratios in LB exhibit increased survival in minimal media. Increased survival could be caused by differential expression of toxin/antitoxin and/or higher metabolic circuit burden within the inoculum, leading to increased escape^44,45^.

Collectively, the results of our screen demonstrate that toxin-inactivator systems cause varying levels of fitness defects when ported into the same genetic context and exhibit a range of toxicities when over-expressed. Inactivator proteins are powerful determinants of this toxicity and therefore represent logical targets for optimizing circuit stability.

### Proteolytic stability of inactivator proteins drives cell killing efficiency

In an environment such as the rhizosphere, *P. fluorescens* may be exposed to heterogenous inducing signals that compromise the efficacy of containment circuits, especially if a high level of toxin is required to overcome the permissive state driven by its cognate inactivator. Therefore, we challenged the killing efficiency of each toxin by measuring liquid growth in the presence of both permissive and non-permissive signals (**Fig. 2A-C**, **Fig. S3**). For comparison, we conducted the same analysis on the toxin circuits lacking an inactivator gene (*tisB* and *ecoRI*). In the absence of an inactivator, inhibition by TisB and EcoRI is a graded response to increasing cumate levels (**Fig. 2A, top).** EcoRI, though, is a more lethal toxin than TisB at lower levels of inducer and, remarkably, exerts no noticeable fitness defect when uninduced, resulting in an almost discrete switch. For dual circuits encoding Type II TA systems, antitoxin expression protects cells from toxin co-expression in an IPTG-dependent manner (**Fig. 2A**). Interestingly, each TA pair exhibits a different steady state “threshold” in our expression system, where antitoxin protection becomes ineffective at a defined level of toxin induction. Antitoxin-mediated protection is much less effective in the *parE-parD* and *relE-relB* systems at lower IPTG levels, which suggests that the high fitness cost of these two circuits (**Fig. 1B,** middle panel) is due to an imbalanced toxin-antitoxin steady state. In contrast to TA systems, colicin E2 and Tse2 are incapable of overcoming the barrier that their respective immunity proteins mount (**Fig. 2A**). Only maximum toxin induction (1 mM cumate) without any immunity induction leads to appreciable cell death by colicin E2. For *tse2-tsi2*, 0.2 mM IPTG is sufficient to protect cells from maximal Tse2 induction. We expect that the relatively short intracellular half-life of antitoxin proteins^46,47^ is crucial for promoting the rapid cell death that occurs with TA systems, whereas the stability of immunity proteins prevents rapid cell death by colicin E2 and Tse2.

**Figure 2.**
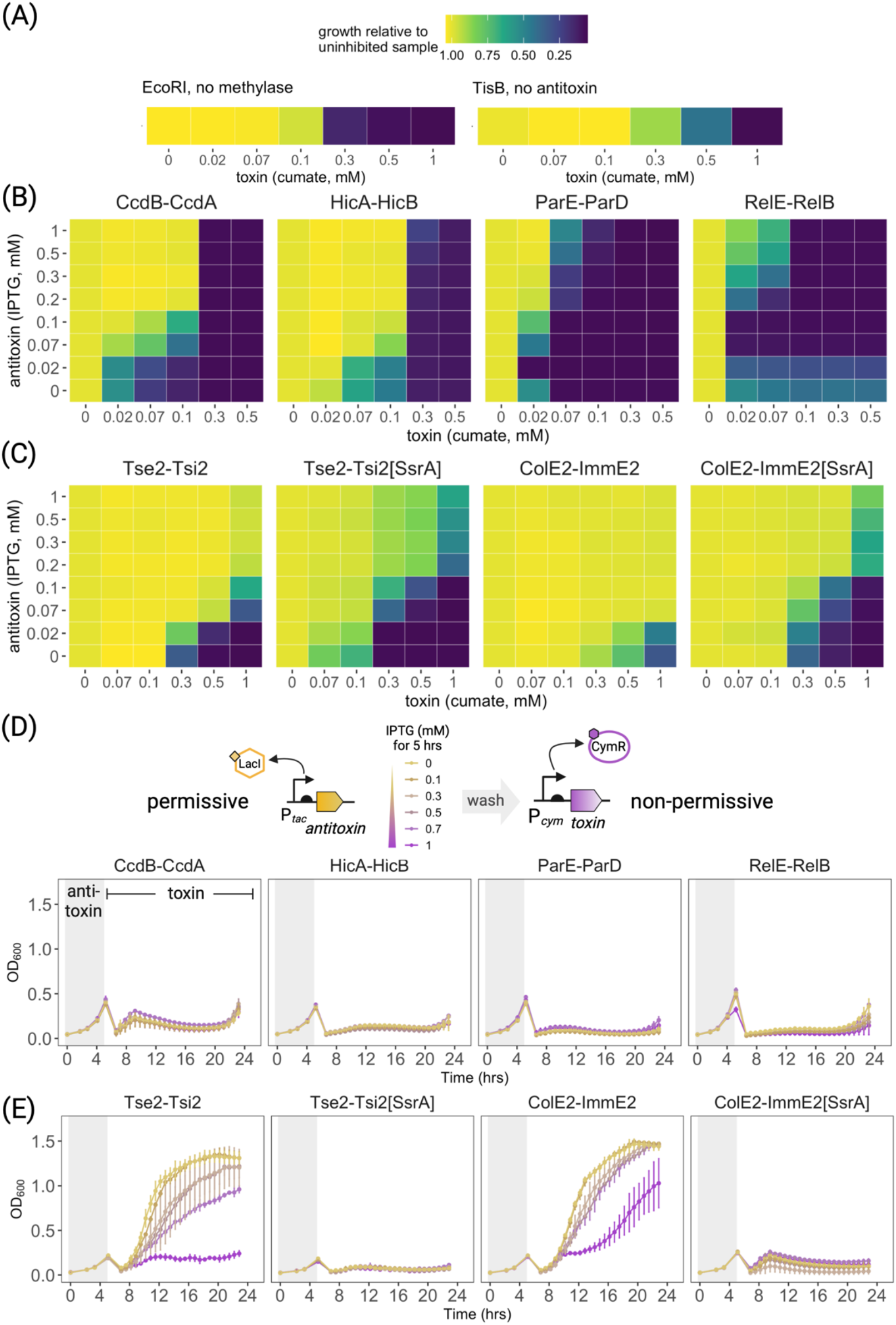
Comparison of co-induction and circuit switching behavior among *P. fluorescens* SBW25 strains with various dual regulatory kill switches. (A-C) Co-induction matrices for *P. fluorescens* carrying (A) *ecoRI* or *tisB* without an inactivator, (B) toxin-antitoxin systems or (C) exotoxin-immunity systems. Cells were grown with the indicated concentrations of one or both inducers. Relative growth is presented (colored boxes), in which the OD_600_ values following 8 hours of induction for each cumate-treated sample is divided by that of the untreated sample at each respective IPTG concentration (i.e., each row). Notably, RelE-RelB samples in (B) without inducers or with 0.02 mM IPTG have a growth defect, making these relative growth values lower than the values at higher IPTG concentrations. (D-E) Growth of (D) toxin-antitoxin expressing strains or (E) exotoxin-immunity expressing strains during a switch from permissive to non-permissive conditions. Strains were grown for five hours with varying levels of IPTG in LB (permissive growth), washed, resuspended in LB with 0.5 mM cumate to an OD_600_ of 0.05, and grown for an additional 16 hours (non-permissive growth). Error bars represent data from three replicates ±SD.

To more directly assess the impact of inactivator stability in shaping kill switch behavior in *P. fluorescens*, we challenged the ability of each toxin-inactivator pair to facilitate containment during a permissive-to-non-permissive switch. Each strain was pre-grown with varying antitoxin induction levels that span the range of P_*tac*_ for five hours, and then challenged with toxin expression for 16 hours. Reassuringly, for TA systems, even maximal antitoxin pre-expression does not impede toxins from driving robust cell death upon induction (**Fig. 2B**). Some growth occurs at the end of the assay (i.e., with *ccdB-ccdA*, *relE-relB*), but is likely due to genetic escape; both strains are fully resistant to toxin induction by the end of the assay (survival ratio of 1; **Fig. S4**). Escape is not surprising considering the increased fitness of inactivated mutants during toxin induction. In contrast, immunity proteins again mount an exceptionally effective barrier against their respective toxins, even without any induction (**Fig. 2B**). These data reveal that there is no level of antitoxin induction from our expression system that enables effective neutralization of fully de-repressed Type II toxin, whereas immunity proteins neutralize their cognate toxins even at basal levels of expression.

Given that the labile antitoxins of TA systems present no apparent barrier to toxin induction (**Fig. 2A-B**), we hypothesized that the longer intracellular half-life of immunity proteins^46,47^ is responsible for increased survival even when toxin is over-expressed. To test this hypothesis, we mimicked the high proteolytic turnover that is a feature of antitoxins^41,42^ in the colicin and Tse2 circuits. To accomplish this, we added the native SBW25 tmRNA sequence (also called an SsrA tag) to the C-terminus of each immunity protein to target it for degradation by ClpXP^48^. To our knowledge, the *sspB*-ClpXP degradation system has not been used for targeted degradation in *P. fluorescens*. Therefore, we selected several SsrA tags than encode modified C-terminal amino acids based on previous studies of SsrA-mediated protein degradation in *E. coli* and *B. subtilis* ^49,50^. C-terminally altered SsrA sequences should recruit the native ClpXP protease with less efficiency, leading to lower rates of turnover compared to the wild type SsrA sequence (-ANDENYGQEFA**LAA**), but still result in lower expression levels of the target protein. Consistent with this expectation, GFP-SsrA[LAA] is completely depleted from cells regardless of inducer level while all four alternative SsrA tags (-DAS, -ILV, -LVC, -AAV) are detectable but at a lower level than the untagged protein (**Fig. S5**). Encouragingly, SsrA-mediated destabilization of the immunity proteins does not cause noticeable toxicity in the absence of inducers but does result in more effective cell death compared to the original circuits when toxin is induced (**Fig. S6**).

We analyzed circuits with immunity-AAV SsrA tags further, as these kill switches are the most potent in *P. fluorescens*. Remarkably, both SsrA-modified toxin-immunity circuits phenocopy Type II TA circuits, exhibiting an improved sensitivity to cumate induction in both inducer assays (**Fig. 2, Fig. S7**) that translates to a 10-100-fold decrease in survival ratio in both rich and minimal media (**Fig. 3**). Altogether, these data show that by reducing the steady state level of immunity protein using targeted proteolysis, we can increase lethality of the *colE2-immE2* and *tse2-tsi2* circuits. This suggests that higher relative proteolytic stability of inactivator proteins can prevent effective on-switching from the immunity-driven off-state. Thus, these data, in combination with the optimal switching behavior of TA systems, indicate that both toxin:inactivator ratios and inactivator turnover rate are critical optimization targets for an effective kill switch.

**Figure 3.**
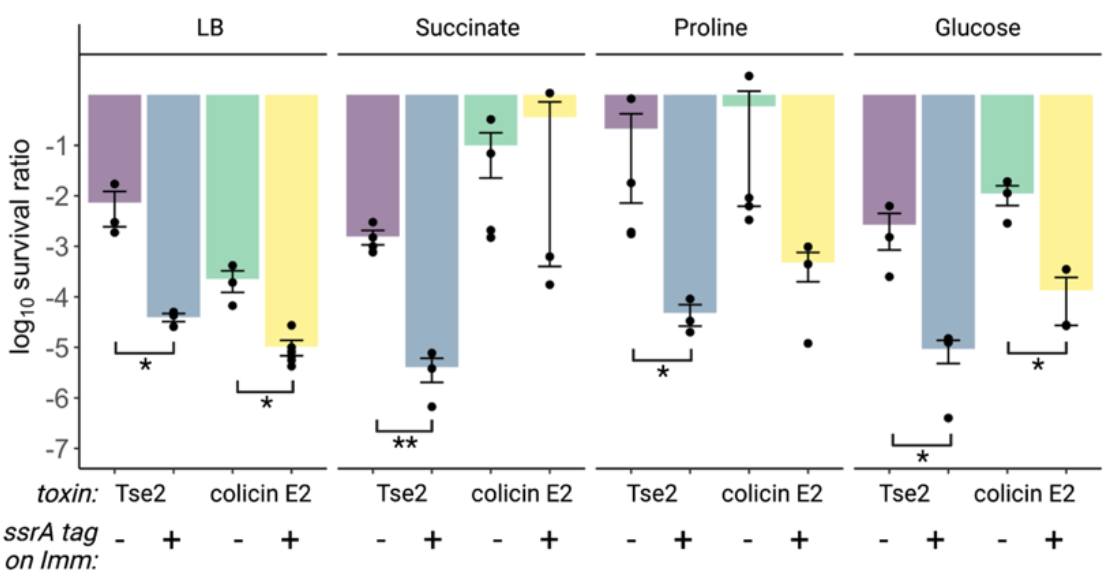
Comparison of survival ratios for *P. fluorescens* SBW25 carrying *tse2-tsi2* and *colE2-immE2* circuits with and without SsrA-modification of the immunity protein. Survival ratio (defined in Fig 1) of *P. fluorescens* SBW25 carrying each toxin-immunity circuit was tested on either rich medium (LB) or minimal M9 media containing 20 mM of the indicated carbon sources. Data represent calculations from three separate replicates ±SEM. Asterisks indicate the following p-value cutoffs obtained from pairwise t-tests performed between the indicated data sets: p>0.05 (none), p<0.05 (one), p<0.01 (two).

### Drivers of mutational escape in *P. fluorescens*

To determine modes of mutational escape in SBW25, we isolated multiple mutants that arose spontaneously on LB agar with cumate (**Fig. 1**) for each toxin circuit and sequenced the entire kill-switch-encoding plasmid to identify causative mutations; 68 such mutants were analyzed in total (**Fig. 4,** Table S4). Altogether, most mutational escape events occurred by re-arrangement of short repeat sequences that resulted in deletions or duplications, ultimately causing frameshifts in circuit components (27 duplications and 12 deletions of 51 total inactivated genes). The majority of mutations in promoter regions were deletions between homologous operator sites (10 of 14 P_*cym*_ mutations), while the remainder were single base substitutions in regulatory regions (3 of 3 LacO mutants, 4 of 14 P_*cym*_ mutants). We suspect that each promoter mutant alters the rate of transcription of the downstream gene to create a permissive level of free toxin. Strikingly, 33 of the 68 mutants we sequenced were inactivated by mutations in *lacI*, which likely causes inactivator over-expression. The majority of *lacI* mutations involve a single direct repeat sequence (TGCCA, beginning at +663, see Table S4) that is well-known as a rearrangement hot spot and a common cause of genetic device failure^15,51,52^. Removal of mutagenic sequences such as these should improve circuit stability. By comparison to repeat-mediated rearrangements, very few point mutations accumulated in protein-coding regions (8 of 51), with all but two resulting in nonsense mutations. The two missense mutations occurred in EcoRI (D133G) and LacI (G252S) and likely cause reduced protein function given the absence of any additional mutations in the circuit. Notably, no mutants contained mobile element insertions in the engineered circuit, consistent with a comprehensive analysis of the SBW25 genome in which no such elements could be identified^53^. A lack of mobile elements has also been reported in the closely related *Pseudomonas protegens* Pf-5^28^.

**Figure 4.**
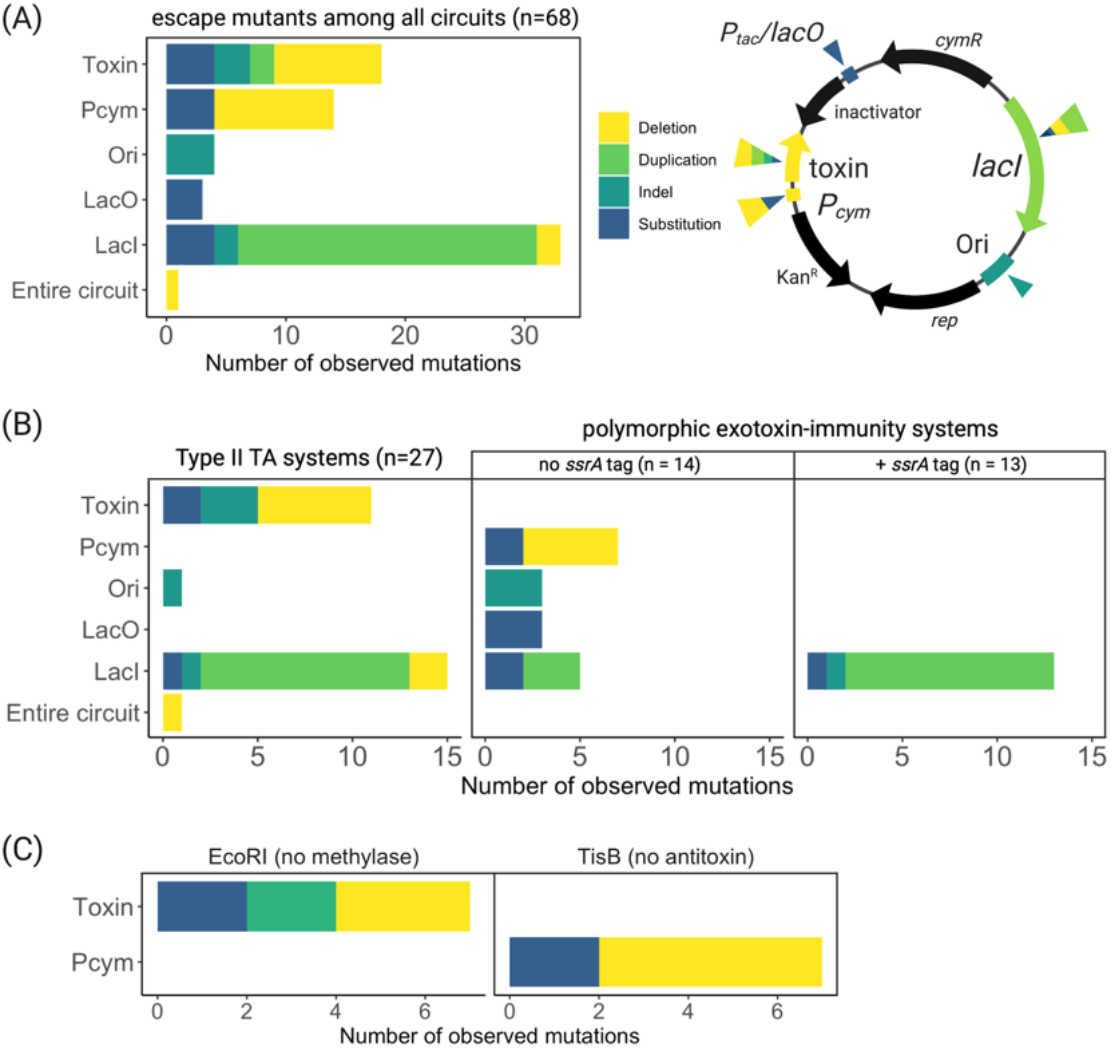
Modes and drivers of kill switch escape in *P. fluorescens* SBW25. Plasmids from seven individual colonies of separate lineages for each escaped strain (except six for *colE2-immE2* and *relE-relB*) were purified and sequenced to determine modes of inactivation. (A) Targets and modes of escape across all sequenced circuits. (B) Spectrum of escape mutations for circuits containing an inactivator gene: Type II TA circuits (left) and *tse2-tsi2* and *colE2-immE2* circuits without (middle) and with (right) SsrA modification of the immunity protein. (C) Spectrum of escape mutations for circuits lacking an inactivator gene, *ecoRI* (left) and *tisB* (right) circuits. For all graphs, n represents the total number of sequenced circuits represented by the data. Notably, some mutations co-occur (e.g., all origin mutations co-occur with at least one other mutation) such that the total number of mutations may exceed the total number of sequenced circuits (see Table S4 for detailed description of the mutations).

For dual circuits encoding an inactivator gene, escape mutations occur either in the toxin gene itself or in regulatory elements controlling expression of the inactivator gene (i.e., *lacI* or P_*tac*_) (**Fig. 4B**). These data indicate that increasing circuit complexity by including an inducible inactivator gene necessarily increases the number of targets for escape mutations. Indeed, computational simulations also find a correlation between circuit complexity (in addition to growth and mutation rates) and the amount of time required for a non-functional mutant to dominate a population of engineered cells^54^. Therefore, circuit complexity should be minimized. For instance, inactivator expression could be controlled with a constitutive promoter to eliminate the need for a second regulator (*lacI*), which was the target of many escape mutations in our analysis. Compared to the TA systems, circuits encoding *colE2-immE2* and *tse2-tsi2* escape by a broader range of mechanisms (P_*cym*_, P_*tac*_, *lacI*), all of which would reduce toxin potency without completely abrogating its activity. Strikingly, in circuits incorporating SsrA tags on the immunity proteins, we find that *lacI* is the sole target of escape mutations (**Fig. 4B**, right). Mutation of *lacI* is also the dominant route of escape for *ccdB-ccdA*, *hicA-hicB*, and *relE-relB* (**Fig. 4B,** left), which share similar survival ratios with the SsrA-tagged immunity systems (~10^-6^). This result implies that de-repression of inactivator genes via *lacI* mutation to continually replenish the inactivator pool is the most viable path to escape for circuits with short-lived inactivator proteins. Moreover, these results also show that by decreasing the protective effect of the inactivator, we can narrow the spectrum of mutational escape to a single locus. This implies that inactivator protection drives selection stringency by decreasing toxicity. In support of this interpretation, experimental evolution studies find that more genetic diversity can be sustained if the strength of selection is lowered^55^. Low selection stringency may also explain the high variability in survival ratios between lineages of SBW25 carrying *colE2-immE2* and *tse2-tsi2* (**Fig. 3**).

For *ecoRI* and *tisB* kill switches that do not include an inactivator gene, we find that promoter mutations inactivated every *tisB* mutant and toxin mutations were present in every *ecoRI* mutant (**Fig. 4C**). These results suggest that the driver of inactivation between *tisB* and *ecoRI* is specific to the toxic modality. Because EcoRI is more potent than TisB at lower levels of cumate (**Fig. 2A**), it is conceivable that reducing expression of TisB is a viable path to escape for this circuit but not for EcoRI. Notably, TisB toxicity is dependent on the formation of tetrameric pores that are sensitive to monomer abundance^56,57^, whereas EcoRI toxicity may be less sensitive to expression level because a single double strand break should be sufficient to inhibit growth. Therefore, drivers of escape may differ by effector class due to intrinsic features of each toxic modality. Altogether, by analyzing inactivating mutations across a wide breadth of toxic effectors from within the same genetic context, we find both commonalities in the mode of escape at the sequence level and drivers of escape that depend on both the toxin and the effectiveness of inactivator protection.

### Long-term stability is inversely correlated with cell killing capacity

Finally, we assessed the relative long-term stability of each engineered circuit under permissive growth conditions, which is an important metric for kill switch performance^20^. We grew each kill switch strain in rich medium over a 10-day period (~100 generations) in a low but sufficient level of IPTG to maintain permissive growth (0.1 mM) and measured survival ratios over time (**Fig. 5**). We find that the least toxic circuits (*tisB*, *colE2-immE2*, *tse2-tsi2*, compare to **Fig. 1B**) are stable over time in their current configuration, with no change in survival ratio over 10 days. This result suggests there is less selective pressure against toxin function in these circuits. In contrast, Type II TA systems and both *colE2-immE2[ssrA]* and *tse2-tsi2[ssrA]* circuits exhibit a daily increase in survival ratio. We anticipate that steadily increasing survival ratios are the result of the preferential accumulation of genetic escape mutants, presumably due to less inactivator protection and thus greater toxicity in the permissive state. Interestingly, EcoRI also drives a constant increase in average survival ratio over 10 days, but at a slower rate than the dual circuits. Accumulation of escape mutants under permissive growth is of great concern in an application with the risk of stochastic toxin induction, which would drive periods of strong selection and compromise containment. Additional optimization of expression or proteolysis would enable a balance between lethal induction and stability over time scales sufficient for the application of interest^13^.

**Figure 4.**
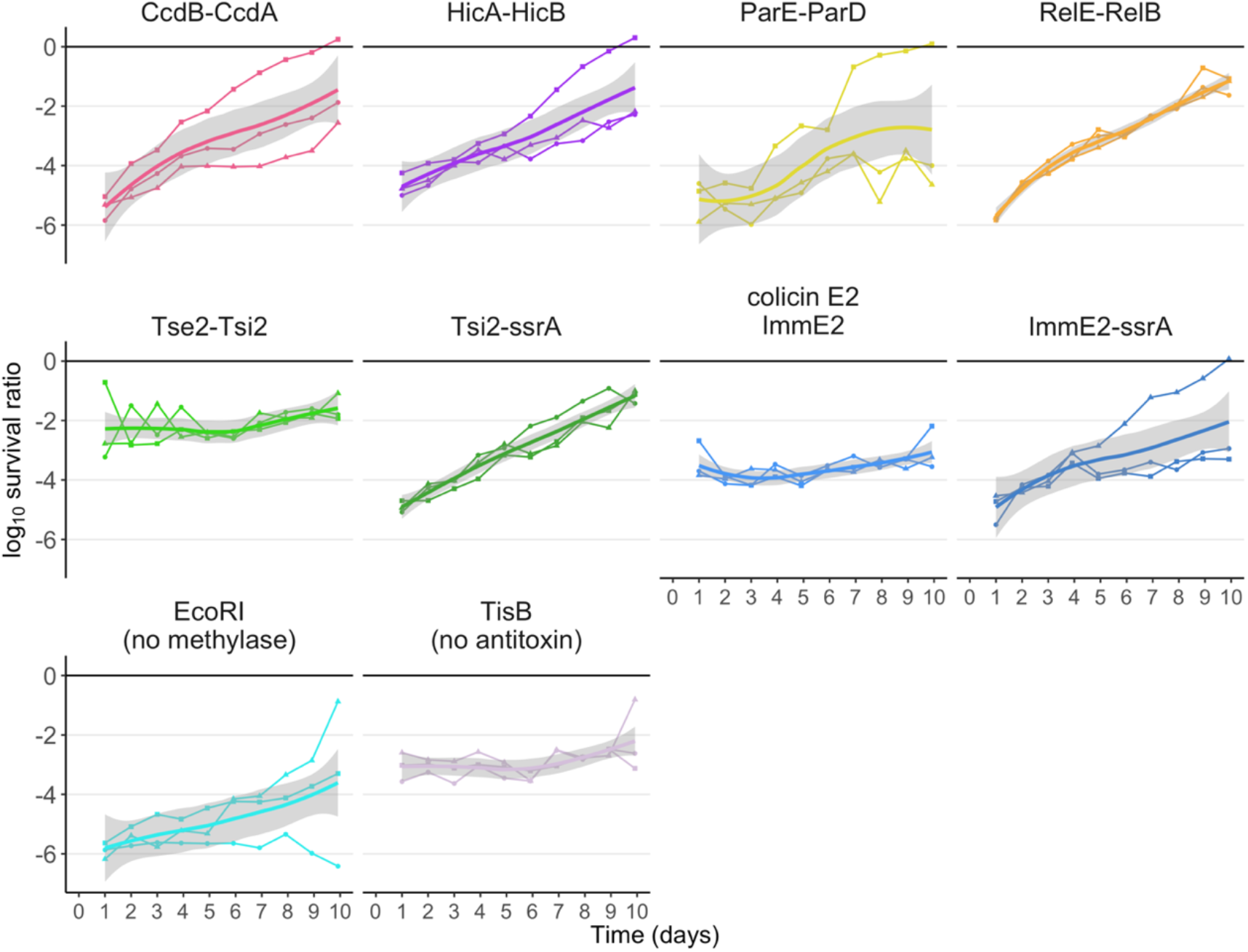
Stability of kill switches during 100 generations of growth. Three independent lineages of each kill switch strain were grown continuously for 10 days in LB supplemented with 0.1 mM IPTG, and survival ratio measured every 24 hours. Individual replicates are shown as colored lines, while the average ±SD is shown as shaded gray regions.

## Conclusions

Here, we present a detailed portfolio of lethality, stability, and mechanisms of escape for kill switches designed with eight different toxin systems in the agriculturally relevant strain *Pseudomonas fluorescens* SBW25. We find that dual circuits with less inactivator protection (TA systems, *colE2-immE2[ssrA]*, *tse2-tsi2[ssrA]*) trigger rapid cell death upon non-permissive induction, but at the cost of decreased long-term genetic stability. In contrast, dual circuits containing highly protective inactivator proteins (*colE2-immE2*, *tse2-tsi2*) are genetically stable over 100 generations, but the potency of toxin induction is dampened. These highly stable systems also take more diverse paths to genetic escape, likely resulting from a lower stringency of genetic selection during non-permissive induction. These results demonstrate the capacity of inactivator proteins to influence kill switch lethality, stability, and escape. Importantly, these results also accentuate the need to focus optimization efforts on achieving a balance between toxin potency (by adjusting toxin-inactivator expression levels) with evolutionary stability (via strategic sequence design) over application lifetimes, an emerging paradigm in synthetic circuit design^20,52^. Finally, our study highlights the importance of testing circuit stability in a host- and condition-specific manner to identify mechanisms of failure relevant to the host strain of interest. In *P. fluorescens*, we find that genetic escape from containment is dominated by rearrangements of small recombinogenic sequences, with no mutants inactivated by mobile elements, in contrast to *E. coli* where mobile element insertions are a common driver of escape^30^. In summary, this indepth analysis of kill switch functionality and stability will guide the design of more stable, highly effective containment systems in agriculturally relevant *P. fluorescens*, by providing new options for toxic effector genes and important considerations for device optimization.

## Methods

### Strains, plasmids, and growth conditions

Primers and gBlocks (Table S3) were ordered from Integrated DNA Technologies (IDT). All gBlock sequences were optimized for *P. putida* using IDT’s online tool, except for *tisB-IstR-2* which was cloned with the wild-type MG1655 sequence (Table S3, NCBI Gene ID 5061525). Ordered gBlocks were PCR-amplified with Phusion High Fidelity polymerase (ThermoFisher) and inserted into pJUMP24-1A^58^ modified with a low copy p15A origin for cloning into *E. coli* via InFusion cloning (Takara Bio), according to the manufacturer’s instructions. LacI/P_tac_ and CymR/P_cym_ were obtained from pMR77 and pMR79^35^ and added to the base vector (pJUMP24-p15A; pTH38) prior to gBlock cloning. Sequences encoding SsrA tags were included in primer sequences and added to *gfp* and both immunity genes during PCR amplification. Primer sequences used to generate InFusion constructs can be supplied upon request. Plasmids were sequence verified by Elim Biopharm. pJUMP24 was maintained in *E. coli* and *P. fluorescens* with 50 mg/mL and 20 mg/mL kanamycin, respectively, in LB broth unless otherwise noted. M9 minimal medium (Sigma) was supplemented with 1 mM MgSO_4_, 100 mM CaCl2 and 20 mM of each carbon source. *P. fluorescens* strains were maintained at 30°C and *E. coli* at 37°C. All overnight cultures were prepared from single colonies plated on solid agar from glycerol stocks and grown in the presence of IPTG, except where indicated.

### Survival ratios, escape analyses, and stability experiments

To measure survival ratios, single colonies of *P. fluorescens* SBW25 or *E. coli* Stellar (Takara) cells on LB- or M9 minimal-agar supplemented with 1 mM IPTG were inoculated into 250 μL of the same liquid medium and grown in a 1.5-mL 96-deep well plate at 240 rpm for ~10 generations. Cells were rinsed and serially diluted, and 5 μL of each ten-fold dilution was plated on the same LB- or M9 minimal-agar containing 1 mM cumate, 1 mM IPTG, or no inducer. CFU/mL were calculated, and the ratio of colonies on cumate relative to total viable colonies on IPTG was calculated as the survival ratio for each strain. For selections in which no colonies were detected on cumate, data is plotted at the limit of detection and noted in the figure caption. Analysis of variance (one way-ANOVA) followed by post-hoc Tukey tests were performed on log-transformed survival ratios for each strain in Figure 1, as a function of medium composition. Pairwise t-tests were performed on log-transformed survival ratios of SsrA-modified and wild type immunity strains from selections in each medium. To identify mutations causing escape, colonies were isolated from separate lineages selected on 1 mM cumate, the plasmid purified (Qiagen), and sequencing performed by Oxford Nanopore (PlasmidSaurus). Each separate lineage came from a single colony grown directly from the original glycerol stock. For stability experiments, strains were grown in 250 μL of LB with 0.1 mM IPTG in a 96-deep well plate as above and passaged with a 1:1000 dilution into fresh medium at 24-hour intervals. Survival ratio measurements were taken at each 24-hour interval for a total of ten days (100 generations).

### Inducer matrix growth curves and permissive-non-permissive switch experiments

All growth curves were conducted in 350-μL, flat-bottom, clear, untreated 96-well polystyrene plates (ThermoFisher) with optical density read in a Synergy H1 (BioTek) at 30°C with linear shaking. To measure the dynamics of each kill switch with competing levels of toxin and antitoxin/immunity, growth curves in 96-well plates were initiated from mid-exponential phase LB cultures without IPTG. Cells were diluted to an initial OD_600_ of 0.05 in 200 μL of LB supplemented with various concentrations of cumate or IPTG and grown for 8 h after which the OD_600_ was measured. To visualize the inducer matrices, relative growth for each treatment of cumate was calculated relative to the sample without any cumate, at each level of IPTG. To measure inhibition, the OD_600_ after 8 h of induction for each cumate-treated sample was normalized to the OD_600_ of the corresponding IPTG only sample to calculate relative growth for each level of antitoxin expression. For permissive-non-permissive switching experiments, overnight cultures of each strain were adjusted to OD_600_ 0.05 in 200 μL of LB supplemented with various concentrations of IPTG (0 to 1 mM), in triplicate. After ~5 h of growth with manual reads taken every hour during growth in a benchtop plate shaker at 1000 rpm, the OD_600_ of each well was re-adjusted to OD_600_ 0.05 in LB and pelleted by centrifugation at 3000 rcf for 4 min. After rinsing 1x in LB, all wells were resuspended in LB supplemented with 0.5 mM cumate, and subsequent reads were taken kinetically every 40 min for at least 16 h. Survival ratios were calculated at the onset of inoculation or after 16 h of growth in cumate as previously described.

## Author Information

^1^Lawrence Livermore National Laboratory, Biosciences and Biotechnology Division, Livermore, CA.

### Author Contributions

T.M.H., D.P.R., D.M.P., Y.J., and M.C.Y. conceived and designed experiments. T.M.H. performed and analyzed experiments. T.M.H. and M.C.Y. wrote the manuscript.

## Acknowledgement

We thank Prof. Christopher Voigt (MIT) for plasmids pMR77 and pMR79. pJUMP24-1A(sfGFP) was a gift from Chris French (Addgene plasmid # 126971). We thank Jenni Chlebek for plasmid pSJC38 (Table S2). We thank Rob Egbert (PNNL) for *P. fluorescens* strain SBW25. We thank Sean Leonard, Jenni Chlebek, and Christina Kang-Yun for helpful discussions. We thank Sean Leonard for extremely critical review of this manuscript. Figures/cartoons were generated by BioRender. This work is supported by the U.S. Department of Energy (DOE), Office of Science, Office of Biological and Environmental Research, Lawrence Livermore National Laboratory SFA “From Sequence to Cell to Population: Secure and Robust Biosystems Design for Environmental Microorganisms.” This work was performed under the auspices of the U.S. DOE by Lawrence Livermore National Laboratory under Contract DE-AC52-07NA27344 (LLNL-JRNL-837127).

## References

(1) Lee, S. Y.; Kim, H. U.; Chae, T. U.; Cho, J. S.; Kim, J. W.; Shin, J. H.; Kim, D. I.; Ko, Y.-S.; Jang, W. D.; Jang, Y.-S. A Comprehensive Metabolic Map for Production of Bio-Based Chemicals. Nat. Catal. 2019, 2 (1), 18–33. https://doi.org/10.1038/s41929-018-0212-4.

(2) Flaiz, M.; Ludwig, G.; Bengelsdorf, F. R.; Dürre, P. Production of the Biocommodities Butanol and Acetone from Methanol with Fluorescent FAST-Tagged Proteins Using Metabolically Engineered Strains of Eubacterium Limosum. Biotechnol. Biofuels 2021, 14 (1), 117. https://doi.org/10.1186/s13068-021-01966-2.

(3) Ye, L.; Lv, X.; Yu, H. Engineering Microbes for Isoprene Production. Metab. Eng. 2016, 38, 125–138. https://doi.org/10.1016/j.ymben.2016.07.005.

(4) Scott, B. M.; Gutiérrez-Vázquez, C.; Sanmarco, L. M.; da Silva Pereira, J. A.; Li, Z.; Plasencia, A.; Hewson, P.; Cox, L. M.; O’Brien, M.; Chen, S. K.; Moraes-Vieira, P. M.; Chang, B. S. W.; Peisajovich, S. G.; Quintana, F. J. Self-Tunable Engineered Yeast Probiotics for the Treatment of Inflammatory Bowel Disease. Nat. Med. 2021, 27 (7), 1212–1222. https://doi.org/10.1038/s41591-021-01390-x.

(5) Riglar, D. T.; Silver, P. A. Engineering Bacteria for Diagnostic and Therapeutic Applications. Nat. Rev. Microbiol. 2018, 16 (4), 214–225. https://doi.org/10.1038/nrmicro.2017.172.

(6) Jaiswal, S.; Shukla, P. Alternative Strategies for Microbial Remediation of Pollutants via Synthetic Biology. Front. Microbiol. 2020, 11, 808. https://doi.org/10.3389/fmicb.2020.00808.

(7) Jia, X.; Li, Y.; Xu, T.; Wu, K. Display of Lead-Binding Proteins on Escherichia Coli Surface for Lead Bioremediation. Biotechnol. Bioeng. 2020, 117 (12), 3820–3834. https://doi.org/10.1002/bit.27525.

(8) Rodríguez, M.; Torres, M.; Blanco, L.; Béjar, V.; Sampedro, I.; Llamas, I. Plant Growth-Promoting Activity and Quorum Quenching-Mediated Biocontrol of Bacterial Phytopathogens by Pseudomonas Segetis Strain P6. Sci. Rep. 2020, 10 (1), 4121. https://doi.org/10.1038/s41598-020-61084-1.

(9) Kristoffersen, P.; Jensen, G. B.; Gerdes, K.; Piskur, J. Bacterial Toxin-Antitoxin Gene System as Containment Control in Yeast Cells. Appl. Environ. Microbiol. 2000, 66 (12), 5524–5526. https://doi.org/10.1128/AEM.66.12.5524-5526.2000.

(10) Chan, C. T. Y.; Lee, J. W.; Cameron, D. E.; Bashor, C. J.; Collins, J. J. “Deadman” and “Passcode” Microbial Kill Switches for Bacterial Containment. Nat. Chem. Biol. 2016, 12 (2), 82–86. https://doi.org/10.1038/nchembio.1979.

(11) Stirling, F.; Naydich, A.; Bramante, J.; Barocio, R.; Certo, M.; Wellington, H.; Redfield, E.; O’Keefe, S.; Gao, S.; Cusolito, A.; Way, J.; Silver, P. Synthetic Cassettes for PH-Mediated Sensing, Counting, and Containment. Cell Rep. 2020, 30 (9), 3139–3148.e4. https://doi.org/10.1016/j.celrep.2020.02.033.

(12) Piraner, D. I.; Abedi, M. H.; Moser, B. A.; Lee-Gosselin, A.; Shapiro, M. G. Tunable Thermal Bioswitches for in Vivo Control of Microbial Therapeutics. Nat. Chem. Biol. 2017, 13 (1), 75–80. https://doi.org/10.1038/nchembio.2233.

(13) Stirling, F.; Bitzan, L.; O’Keefe, S.; Redfield, E.; Oliver, J. W. K.; Way, J.; Silver, P. A. Rational Design of Evolutionarily Stable Microbial Kill Switches. Mol. Cell 2017, 68 (4), 686–697.e3. https://doi.org/10.1016/j.molcel.2017.10.033.

(14) Clark, R. L.; Gordon, G. C.; Bennett, N. R.; Lyu, H.; Root, T. W.; Pfleger, B. F. High-CO2 Requirement as a Mechanism for the Containment of Genetically Modified Cyanobacteria. ACS Synth. Biol. 2018, 7 (2), 384–391. https://doi.org/10.1021/acssynbio.7b00377.

(15) Gallagher, R. R.; Patel, J. R.; Interiano, A. L.; Rovner, A. J.; Isaacs, F. J. Multilayered Genetic Safeguards Limit Growth of Microorganisms to Defined Environments. Nucleic Acids Res. 2015, 43 (3), 1945–1954. https://doi.org/10.1093/nar/gku1378.

(16) Xue, Y.; Qiu, T.; Sun, Z.; Liu, F.; Yu, B. Mercury Bioremediation by Engineered Pseudomonas Putida KT2440 with Adaptationally Optimized Biosecurity Circuit. Environ. Microbiol. n/a (n/a). https://doi.org/10.1111/1462-2920.16038.

(17) Jensen, L. B.; Ramos, J. L.; Kaneva, Z.; Molin, S. A Substrate-Dependent Biological Containment System for Pseudomonas Putida Based on the Escherichia Coli Gef Gene. Appl. Environ. Microbiol. 1993, 59 (11), 3713–3717. https://doi.org/10.1128/aem.59.11.3713-3717.1993.

(18) Rottinghaus, A. G.; Ferreiro, A.; Fishbein, S. R. S.; Dantas, G.; Moon, T. S. Genetically Stable CRISPR-Based Kill Switches for Engineered Microbes. Nat. Commun. 2022, 13 (1), 672. https://doi.org/10.1038/s41467-022-28163-5.

(19) Munthali, M. T.; Timmis, K. N.; Diaz, E. Use of Colicin E3 for Biological Containment of Microorganisms. Appl. Environ. Microbiol. 1996, 62 (5), 1805–1807. https://doi.org/10.1128/aem.62.5.1805-1807.1996.

(20) Stirling, F.; Silver, P. A. Controlling the Implementation of Transgenic Microbes: Are We Ready for What Synthetic Biology Has to Offer? Mol. Cell 2020, 78 (4), 614–623. https://doi.org/10.1016/j.molcel.2020.03.034.

(21) Ahrenholtz, I.; Lorenz, M. G.; Wackernagel, W. A Conditional Suicide System in Escherichia Coli Based on the Intracellular Degradation of DNA. Appl. Environ. Microbiol. 1994, 60 (10), 3746–3751. https://doi.org/10.1128/aem.60.10.3746-3751.1994.

(22) Callura, J. M.; Dwyer, D. J.; Isaacs, F. J.; Cantor, C. R.; Collins, J. J. Tracking, Tuning, and Terminating Microbial Physiology Using Synthetic Riboregulators. Proc. Natl. Acad. Sci. 2010, 107 (36), 15898–15903. https://doi.org/10.1073/pnas.1009747107.

(23) Balan, A.; Schenberg, A. C. G. A Conditional Suicide System for Saccharomyces Cerevisiae Relying on the Intracellular Production of the Serratia Marcescens Nuclease. Yeast 2005, 22 (3), 203–212. https://doi.org/10.1002/yea.1203.

(24) Ronchel, M. C.; Molina, L.; Witte, A.; Lutbiz, W.; Molin, S.; Ramos, J. L.; Ramos, C. Characterization of Cell Lysis in Pseudomonas Putida Induced upon Expression of Heterologous Killing Genes. Appl. Environ. Microbiol. 1998, 64 (12), 4904–4911. https://doi.org/10.1128/AEM.64.12.4904-4911.1998.

(25) Bej, A. K.; Molin, S.; Perlin, M.; Atlas, R. M. Maintenance and Killing Efficiency of Conditional Lethal Constructs in Pseudomonas Putida. J. Ind. Microbiol. 1992, 10 (2), 79–85. https://doi.org/10.1007/BF01583839.

(26) Moser, F.; Broers, N. J.; Hartmans, S.; Tamsir, A.; Kerkman, R.; Roubos, J. A.; Bovenberg, R.; Voigt, C. A. Genetic Circuit Performance under Conditions Relevant for Industrial Bioreactors. ACS Synth. Biol. 2012, 1 (11), 555–564. https://doi.org/10.1021/sb3000832.

(27) Ishikawa, M.; Kojima, T.; Hori, K. Development of a Biocontained Toluene-Degrading Bacterium for Environmental Protection. Microbiol. Spectr. 9 (1), e00259–21. https://doi.org/10.1128/Spectrum.00259-21.

(28) Mavrodi, D. V.; Loper, J. E.; Paulsen, I. T.; Thomashow, L. S. Mobile Genetic Elements in the Genome of the Beneficial Rhizobacterium Pseudomonas Fluorescens Pf-5. BMC Microbiol. 2009, 9 (1), 8. https://doi.org/10.1186/1471-2180-9-8.

(29) Csörgő, B.; Fehér, T.; Tímár, E.; Blattner, F. R.; Pósfai, G. Low-Mutation-Rate, Reduced-Genome Escherichia Coli: An Improved Host for Faithful Maintenance of Engineered Genetic Constructs. Microb. Cell Factories 2012, 11 (1), 11. https://doi.org/10.1186/1475-2859-11-11.

(30) Consuegra, J.; Gaffé, J.; Lenski, R. E.; Hindré, T.; Barrick, J. E.; Tenaillon, O.; Schneider, D. Insertion-Sequence-Mediated Mutations Both Promote and Constrain Evolvability during a Long-Term Experiment with Bacteria. Nat. Commun. 2021, 12 (1), 980. https://doi.org/10.1038/s41467-021-21210-7.

(31) Ganeshan, G.; Manoj Kumar, A. Pseudomonas Fluorescens, a Potential Bacterial Antagonist to Control Plant Diseases. J. Plant Interact. 2005, 1 (3), 123–134. https://doi.org/10.1080/17429140600907043.

(32) Setten, L.; Soto, G.; Mozzicafreddo, M.; Fox, A. R.; Lisi, C.; Cuccioloni, M.; Angeletti, M.; Pagano, E.; Díaz-Paleo, A.; Ayub, N. D. Engineering Pseudomonas Protegens Pf-5 for Nitrogen Fixation and Its Application to Improve Plant Growth under Nitrogen-Deficient Conditions. PLOS ONE 2013, 8 (5), e63666. https://doi.org/10.1371/journal.pone.0063666.

(33) Hernández-León, R.; Rojas-Solís, D.; Contreras-Pérez, M.; Orozco-Mosqueda, Ma. del C.; Macías-Rodríguez, L. I.; Reyes-de la Cruz, H.; Valencia-Cantero, E.; Santoyo, G. Characterization of the Antifungal and Plant Growth-Promoting Effects of Diffusible and Volatile Organic Compounds Produced by Pseudomonas Fluorescens Strains. Biol. Control 2015, 81, 83–92. https://doi.org/10.1016/j.biocontrol.2014.11.011.

(34) Haskett, T. L.; Tkacz, A.; Poole, P. S. Engineering Rhizobacteria for Sustainable Agriculture. ISME J. 2021, 15 (4), 949–964. https://doi.org/10.1038/s41396-020-00835-4.

(35) Ryu, M.-H.; Zhang, J.; Toth, T.; Khokhani, D.; Geddes, B. A.; Mus, F.; Garcia-Costas, A.; Peters, J. W.; Poole, P. S.; Ané, J.-M.; Voigt, C. A. Control of Nitrogen Fixation in Bacteria That Associate with Cereals. Nat. Microbiol. 2020, 5 (2), 314–330. https://doi.org/10.1038/s41564-019-0631-2.

(36) Kamilova, F.; Kravchenko, L. V.; Shaposhnikov, A. I.; Azarova, T.; Makarova, N.; Lugtenberg, B. Organic Acids, Sugars, and l-Tryptophane in Exudates of Vegetables Growing on Stonewool and Their Effects on Activities of Rhizosphere Bacteria. Mol. Plant-Microbe Interactions® 2006, 19 (3), 250–256. https://doi.org/10.1094/MPMI-19-0250.

(37) Liu, S.; Hu, X.; Lohrke, S. M.; Baker, C. J.; Buyer, J. S.; de Souza, J. T.; Roberts, D. P. Y. 2007. Role of SdhA and PfkA and Catabolism of Reduced Carbon during Colonization of Cucumber Roots by Enterobacter Cloacae. Microbiology 153 (9), 3196–3209. https://doi.org/10.1099/mic.0.2006/005538-0.

(38) Cascales, E.; Buchanan, S. K.; Duché, D.; Kleanthous, C.; Lloubès, R.; Postle, K.; Riley, M.; Slatin, S.; Cavard, D. Colicin Biology. Microbiol. Mol. Biol. Rev. 2007, 71 (1), 158–229. https://doi.org/10.1128/MMBR.00036-06.

(39) Hood, R. D.; Singh, P.; Hsu, F.; Güvener, T.; Carl, M. A.; Trinidad, R. R. S.; Silverman, J. M.; Ohlson, B. B.; Hicks, K. G.; Plemel, R. L.; Li, M.; Schwarz, S.; Wang, W. Y.; Merz, A. J.; Goodlett, D. R.; Mougous, J. D. A Type VI Secretion System of Pseudomonas Aeruginosa Targets a Toxin to Bacteria. Cell Host Microbe 2010, 7 (1), 25–37. https://doi.org/10.1016/j.chom.2009.12.007.

(40) James, R.; Kleanthous, C.; Moore, G. R. Y. 1996. The Biology of E Colicins: Paradigms and Paradoxes. Microbiology 142 (7), 1569–1580. https://doi.org/10.1099/13500872-142-7-1569.

(41) Van Melderen, L.; Bernard, P.; Couturier, M. Lon-Dependent Proteolysis of CcdA Is the Key Control for Activation of CcdB in Plasmid-Free Segregant Bacteria. Mol. Microbiol. 1994, 11 (6), 1151–1157. https://doi.org/10.1111/j.1365-2958.1994.tb00391.x.

(42) Hansen, S.; Vulić, M.; Min, J.; Yen, T.-J.; Schumacher, M. A.; Brennan, R. G.; Lewis, K. Regulation of the Escherichia Coli HipBA Toxin-Antitoxin System by Proteolysis. PLOS ONE 2012, 7 (6), e39185. https://doi.org/10.1371/journal.pone.0039185.

(43) Ho, B. T.; Dong, T. G.; Mekalanos, J. J. A View to a Kill: The Bacterial Type VI Secretion System. Cell Host Microbe 2014, 15 (1), 9–21. https://doi.org/10.1016/j.chom.2013.11.008.

(44) Rugbjerg, P.; Sommer, M. O. A. Overcoming Genetic Heterogeneity in Industrial Fermentations. Nat. Biotechnol. 2019, 37 (8), 869–876. https://doi.org/10.1038/s41587-019-0171-6.

(45) Carneiro, S.; Ferreira, E. C.; Rocha, I. Metabolic Responses to Recombinant Bioprocesses in Escherichia Coli. J. Biotechnol. 2013, 164 (3), 396–408. https://doi.org/10.1016/j.jbiotec.2012.08.026.

(46) Wojdyla, J. A.; Fleishman, S. J.; Baker, D.; Kleanthous, C. Structure of the Ultra-High-Affinity Colicin E2 DNase–Im2 Complex. J. Mol. Biol. 2012, 417 (1), 79–94. https://doi.org/10.1016/j.jmb.2012.01.019.

(47) Robb, C. S.; Robb, M.; Nano, F. E.; Boraston, A. B. The Structure of the Toxin and Type Six Secretion System Substrate Tse2 in Complex with Its Immunity Protein. Structure 2016, 24 (2), 277–284. https://doi.org/10.1016/j.str.2015.11.012.

(48) Janssen, B. D.; Hayes, C. S. The TmRNA Ribosome Rescue System. Adv. Protein Chem. Struct. Biol. 2012, 86, 151–191. https://doi.org/10.1016/B978-0-12-386497-0.00005-0.

(49) Griffith, K. L.; Grossman, A. D. Inducible Protein Degradation in Bacillus Subtilis Using Heterologous Peptide Tags and Adaptor Proteins to Target Substrates to the Protease ClpXP. Mol. Microbiol. 2008, 70 (4), 1012–1025. https://doi.org/10.1111/j.1365-2958.2008.06467.x.

(50) Karzai, A. W.; Roche, E. D.; Sauer, R. T. The SsrA–SmpB System for Protein Tagging, Directed Degradation and Ribosome Rescue. Nat. Struct. Biol. 2000, 7 (6), 449–455. https://doi.org/10.1038/75843.

(51) Farabaugh, P. J.; Schmeissner, U.; Hofer, M.; Miller, J. H. Genetic Studies of the Lac Repressor: VII. On the Molecular Nature of Spontaneous Hotspots in the LacI Gene of Escherichia Coli. J. Mol. Biol. 1978, 126 (4), 847–863. https://doi.org/10.1016/0022-2836(78)90023-2.

(52) Renda, B. A.; Hammerling, M. J.; Barrick, J. E. Engineering Reduced Evolutionary Potential for Synthetic Biology. Mol. Biosyst. 2014, 10 (7), 1668–1678. https://doi.org/10.1039/c3mb70606k.

(53) Silby, M. W.; Cerdeño-Tárraga, A. M.; Vernikos, G. S.; Giddens, S. R.; Jackson, R. W.; Preston, G. M.; Zhang, X.-X.; Moon, C. D.; Gehrig, S. M.; Godfrey, S. A.; Knight, C. G.; Malone, J. G.; Robinson, Z.; Spiers, A. J.; Harris, S.; Challis, G. L.; Yaxley, A. M.; Harris, D.; Seeger, K.; Murphy, L.; Rutter, S.; Squares, R.; Quail, M. A.; Saunders, E.; Mavromatis, K.; Brettin, T. S.; Bentley, S. D.; Hothersall, J.; Stephens, E.; Thomas, C. M.; Parkhill, J.; Levy, S. B.; Rainey, P. B.; Thomson, N. R. Genomic and Genetic Analyses of Diversity and Plant Interactions of Pseudomonas Fluorescens. Genome Biol. 2009, 10 (5), R51. https://doi.org/10.1186/gb-2009-10-5-r51.

(54) Arkin, A. P.; Fletcher, D. A. Fast, Cheap and Somewhat in Control. Genome Biol. 2006, 7 (8), 114. https://doi.org/10.1186/gb-2006-7-8-114.

(55) Barrick, J. E.; Lenski, R. E. Genome Dynamics during Experimental Evolution. Nat. Rev. Genet. 2013, 14 (12), 827–839. https://doi.org/10.1038/nrg3564.

(56) Schneider, V.; Wadhwani, P.; Reichert, J.; Bürck, J.; Elstner, M.; Ulrich, A. S.; Kubař, T. Tetrameric Charge-Zipper Assembly of the TisB Peptide in Membranes-Computer Simulation and Experiment. J. Phys. Chem. B 2019, 123 (8), 1770–1779. https://doi.org/10.1021/acs.jpcb.8b12087.

(57) Wilmaerts, D.; Dewachter, L.; De Loose, P.-J.; Bollen, C.; Verstraeten, N.; Michiels, J. HokB Monomerization and Membrane Repolarization Control Persister Awakening. Mol. Cell 2019, 75 (5), 1031–1042.e4. https://doi.org/10.1016/j.molcel.2019.06.015.

(58) Valenzuela-Ortega, M.; French, C. Joint Universal Modular Plasmids (JUMP): A Flexible Vector Platform for Synthetic Biology. Synth. Biol. 2021, 6 (1), ysab003. https://doi.org/10.1093/synbio/ysab003.

