## Supplemental information (Figures S1-S7, Tables S1-S4) for "Comprehensive analysis of kill switch toxins in plant-beneficial *Pseudomonas fluorescens* reveals drivers of lethality, stability, and escape"

***Supporting Figures*** 2

Figure S1. Fold change of $P_{cym}$ and $P_{tac}$ promoters in the dual-regulatory plasmid in *P. fluorescens* SBW25 2

Figure S2. Survival ratios of *P. fluorescens* SBW25 kill switch strains from inoculum cultures grown with or without inactivator over-expression by IPTG. 3

Figure S3. Co-induction matrices of Type II toxin-antitoxin kill switches in SBW25 4

Figure S4. Survival ratios of SBW25 strains with TA systems before and after a permissive to non-permissive switch in liquid media 5

Figure S5. SsrA (tmRNA) degradation tags increase turnover of GFP in *P. fluorescens* 6

Figure S6. Cumate induction of SBW25 carrying *tse2-tsi2* or *colE2-immE2* circuits with various *ssrA* modifications of the immunity gene in liquid LB 7

Figure S7. Co-induction matrices of colicin-Immunity E2 and Tse2-Tsi2 kill switches with and without SsrA immunity modification in SBW25 8

***Supporting Tables*** 9

Table S1. Toxin-inactivator systems used in this study 9

Table S2. Strains and plasmids used in this study. 10

Table S3. gBlock sequences of kill switch circuits. 12

Table S4. Escape mutations for all circuits. 16

#### Supporting Figures

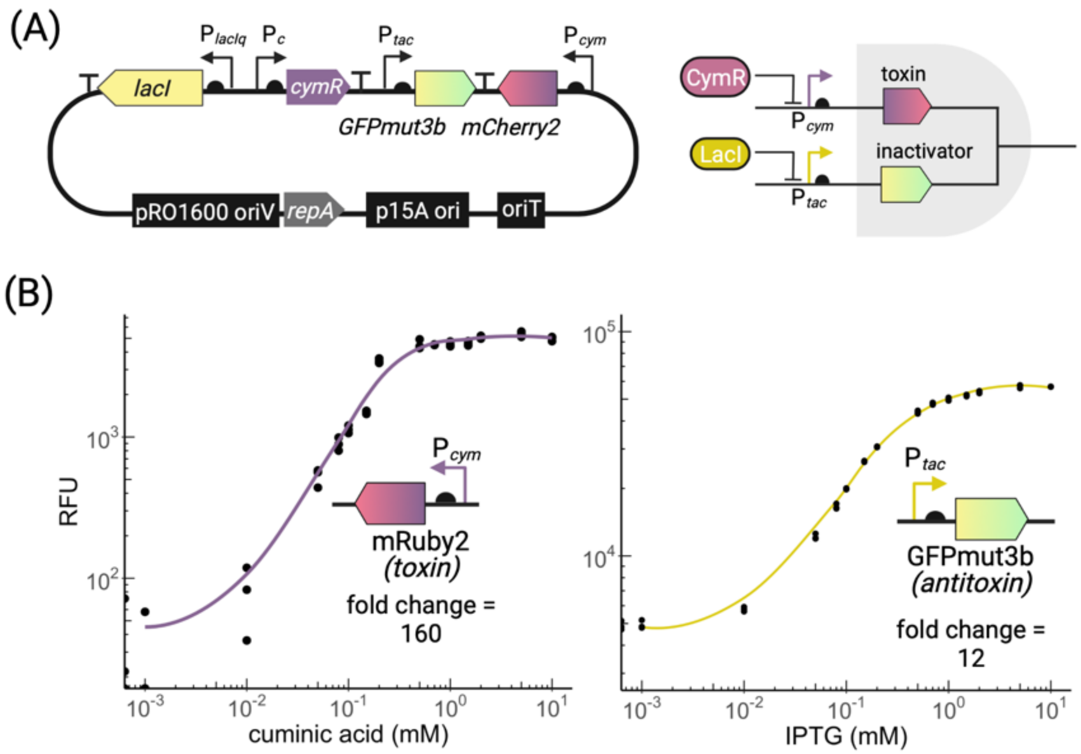


**Figure S1. Fold change of $P_{cym}$ and $P_{tac}$ promoters in the dual-regulatory plasmid in *P. fluorescens* SBW25.** (A) Schematic of plasmid and circuit logic. (B) Fold change of $P_{cym}$ using mRuby2 fluorescence and of $P_{tac}$ using GFPmut3b fluorescence. Cultures were induced in triplicate from mid-log. Fluorescence was measured after 6 hours and calculated relative to $OD_{600}$ at each time point. Individual replicate values and a trendline are shown.

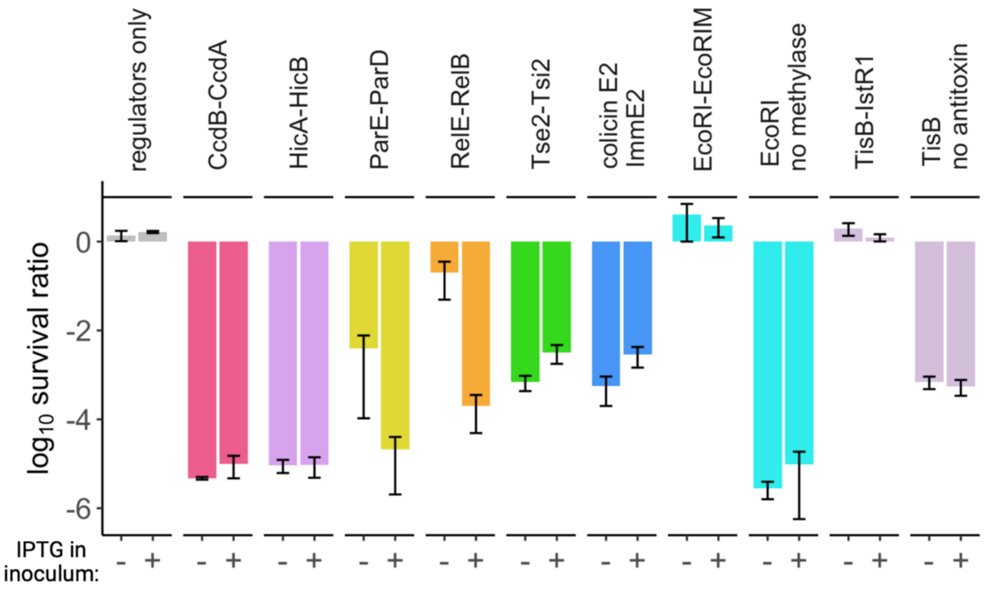


**Figure S2. Survival ratios of *P. fluorescens* SBW25 kill switch strains from inoculum cultures grown with or without inactivator over-expression by IPTG.** Survival ratio is the number of viable cell counts on non-permissive medium (LB with 1 mM cumate) relative to the corresponding CFU/mL on permissive media (LB with 1 mM IPTG). Data is presented as the average of three individual replicates ±SEM.

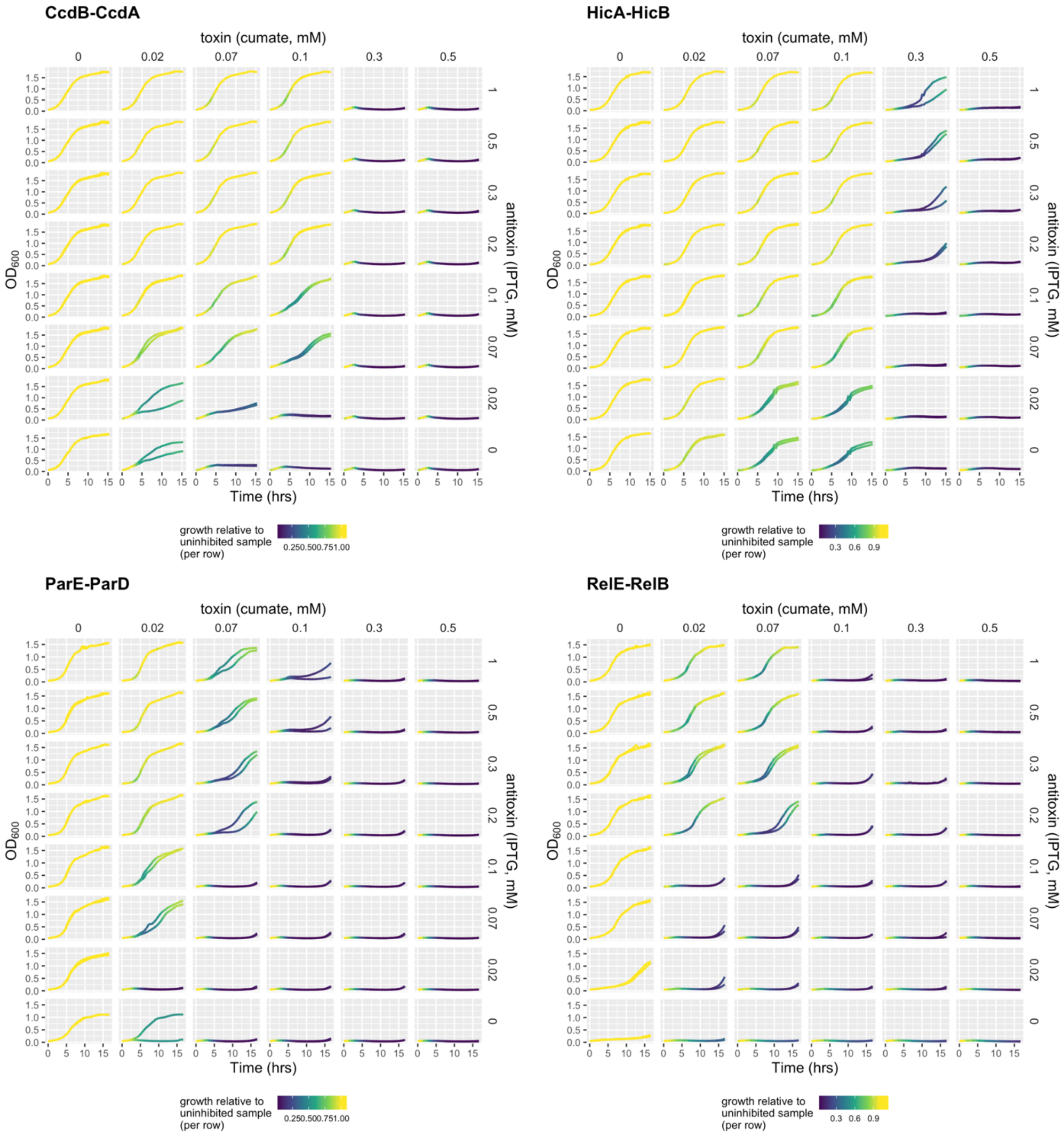


**Figure S3. Co-induction matrices of Type II toxin-antitoxin kill switches in SBW25.** Data from the 8-hour time point are presented in Fig. 2B. The full growth curve is plotted and colored with a heatmap of relative culture density calculated as in Fig. 2: the $OD_{600}$ value of each cumate-treated sample is measured as a fraction of the untreated sample in each IPTG treatment group (i.e., relative growth of each cumate-treated culture is calculated per-row). The relatively high fitness cost of uninduced RelE and ParE is evident in the lower left square of each plot (samples with no inducers added). Two replicates were performed for each treatment, which are plotted separately.

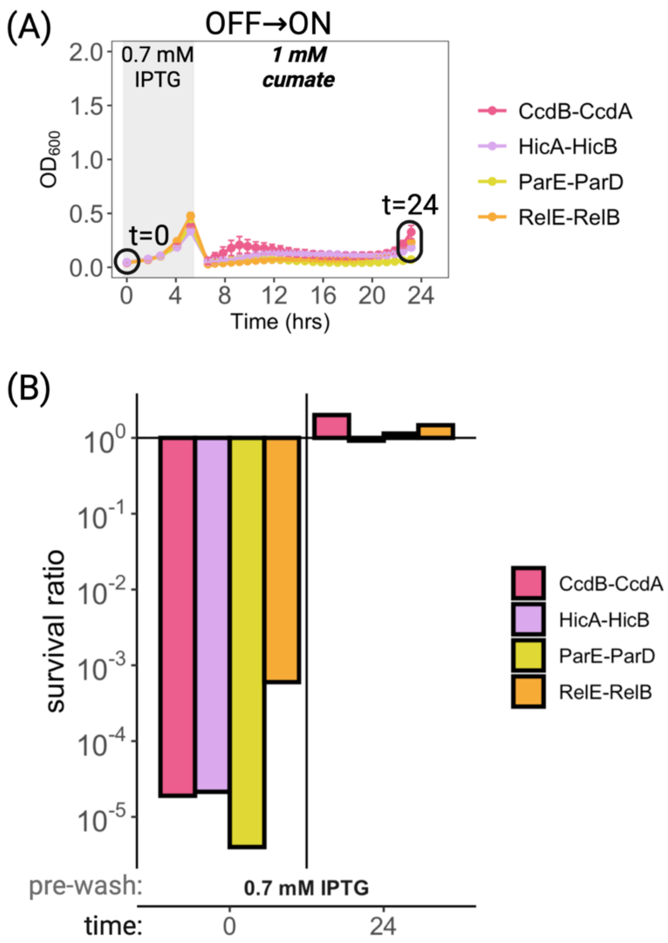


**Figure S4. Survival ratios of SBW25 strains with TA systems before and after a permissive to non-permissive switch in liquid media.** (A) Growth curve as in Figure 2A with 0.7 mM IPTG added prior to switching to 1 mM cumate after a wash step. (B) Survival ratios of each culture from (A) at the onset of the growth curve in permissive conditions (t=0) and at the end following a switch into non-permissive conditions (t=24). All samples treated with 1 mM cumate are insensitive to cumate induction by the end of the assay. Survival ratio is number of viable cell counts on non-permissive medium (LB with 1 mM cumate) relative to the corresponding CFU/mL on permissive media (LB with 1 mM IPTG).

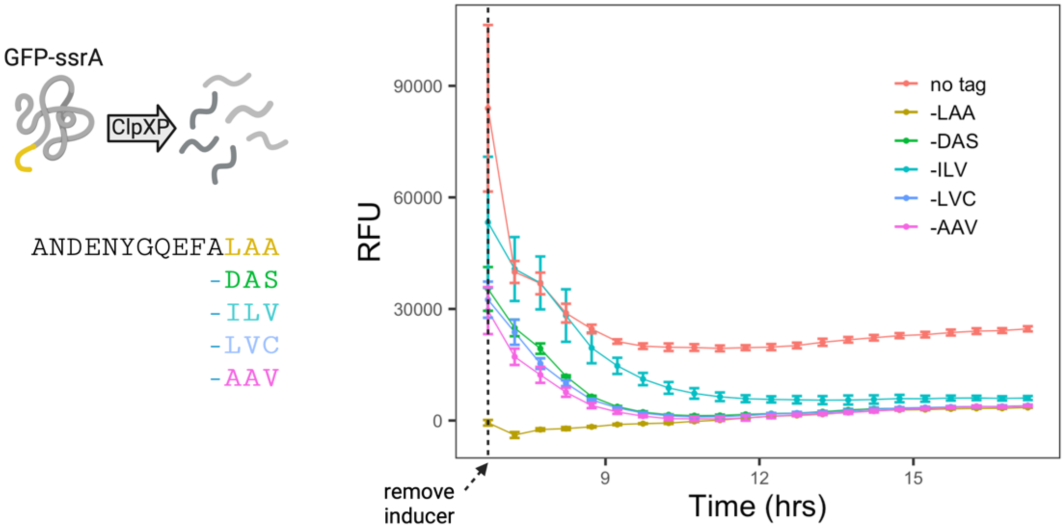


**Figure S5. SsrA** (tmRNA) **degradation tags increase turnover of GFP in *P. fluorescens*.** Left, top: Addition of the SsrA sequence (yellow) onto GFP targets it for degradation by the native ClpXP protease. Left, bottom: SsrA sequences tested in this study. The wild type SsrA sequence includes the conserved -LAA ClpXP binding site. Right: Decay of GFP fluorescence over time following removal of IPTG inducer from SBW25 cells expressing GFP-SsrA variants from $P_{tac}$. RFU (relative fluorescence units) is plotted as absolute GFP fluorescence relative to $OD_{600}$ of each culture over time. The wild type -LAA tag and leads to complete depletion of GFP in SBW25 even under 1 mM IPTG $P_{tac}$ induction. Data are the average ±SD for three independent replicates.

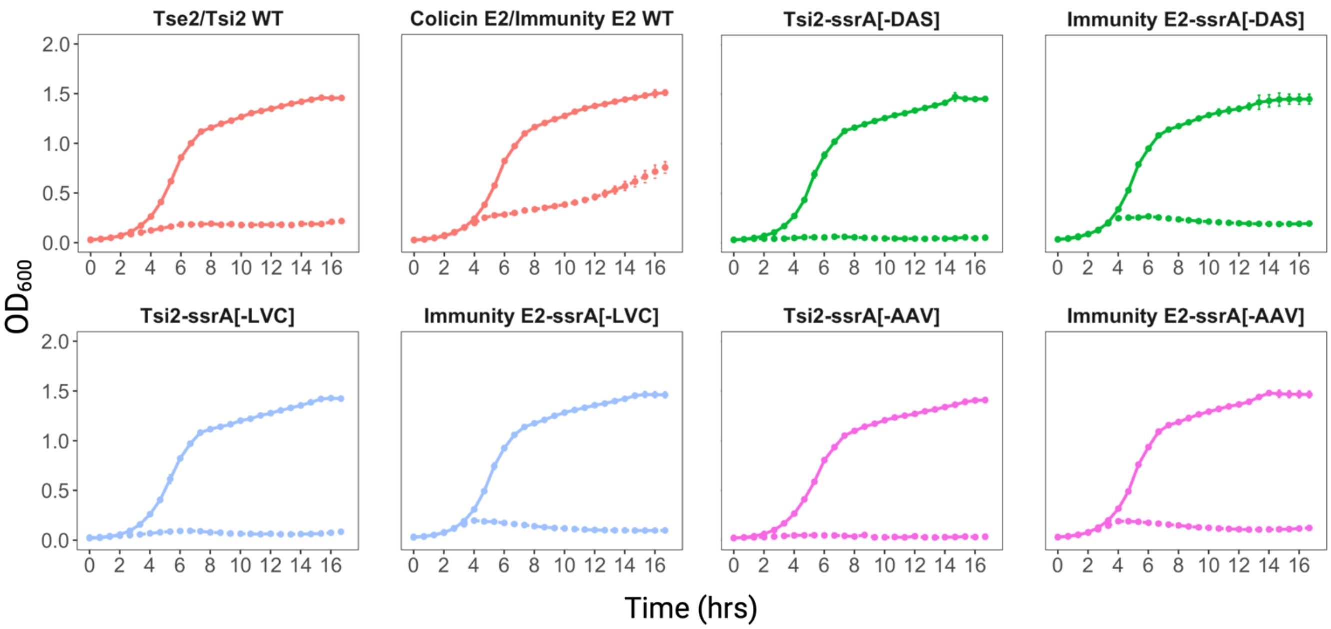


**Figure S6. Cumate induction of SBW25 carrying *tse2-tsi2* or *colE2-immE2* circuits with various *ssrA* modifications of the immunity gene in liquid LB.** Solid lines: growth in the absence of inducers. Hatched lines: 0.5 mM cumate added to induce toxin expression. Data are the average ±SD for three independent replicates.

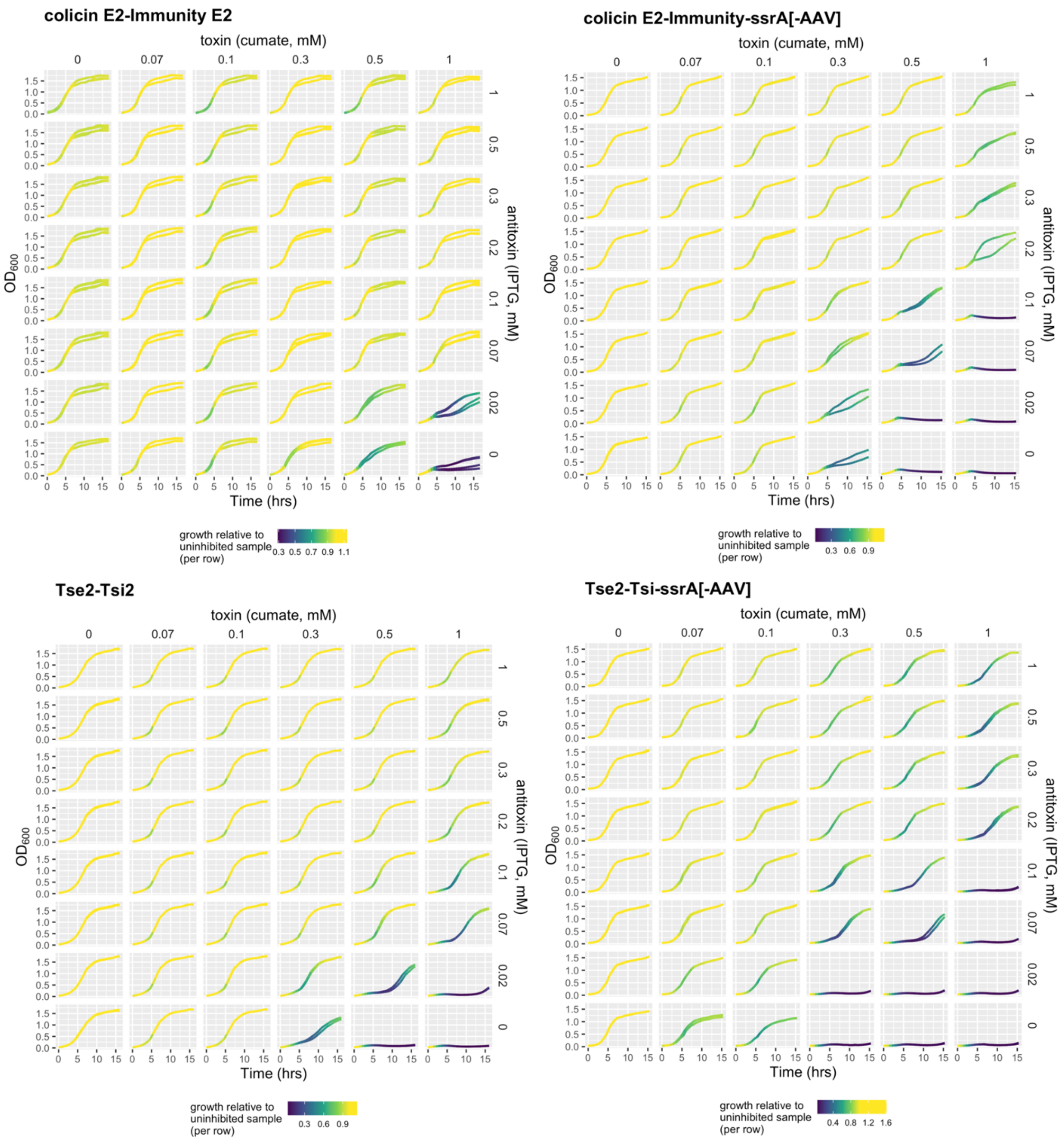


**Figure S7. Co-induction matrices of colicin-Immunity E2 and Tse2-Tsi2 kill switches with and without SsrA immunity modification in SBW25.** Growth curves are colored by relative growth as defined in Fig. S5. Individual replicates are plotted separately.

### Supporting Tables

**Table S1. Toxin-inactivator systems used in this study**

| Toxin Family | Toxin | Inactivator | Activity | Origin |
|---|---|---|---|---|
| Type I TA | TisB | IstR sRNA | Ionophore | *E. coli* K-12 |
| Type II TA | CcdB<br>ParE<br>HicA<br>RelE | CcdA<br>ParD<br>HicB<br>RelB | Gyrase inhibitor<br>Gyrase inhibitor<br>Endoribonuclease<br>Endoribonuclease | F-plasmid<br>Plasmid RK2<br>*E. coli* K-12<br>*E. coli* K-12 |
| Colicin/ Bacteriocin | Colicin E2 | Immunity E2 | Nonspecific DNase | Plasmid ColE2 |
| T6SS | Tse2 | Tsi2 | Cytosolic NAD-dependent predicted ADP-ribosylase. | *Pseudomonas aeruginosa* PAO1 |
| Restriction Endonuclease | EcoRIR | EcoRIM | Endoribonuclease: cleaves DNA at G↓AATTC | *E. coli* |

**Table S2. Strains and plasmids used in this study.**

| Strain | Description | Source |
|---|---|---|
| *Pseudomonas fluorescens* SBW25 | Wild type | Rob Egbert, PNNL |
| *E. coli* HST08 | F-, endA1, supE44, thi-1, recA1, relA1, gyrA96, phoA, Φ80d lacZΔ M15, Δ(lacZYA-argF) U169, Δ(mrr-hsdRMS-mcrBC), ΔmcrA, λ- | Takara Bio |
| **Plasmid** | **Description** | **Source** |
| pTH38 | Derivative of pJUMP24 with p15A ori for lower copy number replication in *E. coli.* | This study |
| pTH48 | pJUMP24-p15A (pTH38) with constitutively expressed LacI and CymR. | This study |
| pTH51 | Kill switch reporter construct. pJUMP24-p15A constitutively expressing LacI and CymR (pTH48) with $P_{tac}$GFP and $P_{cym}$mRuby2 for measuring promoter activity. | This study |
| pTH52 | CcdB-CcdA kill switch. pJUMP24-p15A constitutively expressing LacI and CymR (pTH48) with $P_{tac}$*ccdA* and $P_{cym}$*ccdB* encoded. | This study |
| pTH80 | HicA-HicB kill switch. Constitutive expression of LacI and CymR, inducible $P_{tac}$*hicB* and $P_{cym}$*hicA.* | This study |
| pTH81 | EcoRI-EcoRIM kill switch. Constitutive expression of LacI and CymR, inducible $P_{tac}$*ecoRIM* and $P_{cym}$*ecoRI.* | This study |
| pTH82 | ParE-ParD kill switch. Constitutive expression of LacI and CymR, inducible $P_{tac}$*parD* and $P_{cym}$*parE.* | This study |
| pSJC38 | RelE-RelB kill switch. Constitutive expression of LacI and CymR, inducible $P_{tac}$*relB* and $P_{cym}$*relE.* | This study |
| pTH107 | Colicin E2-immunity E2 kill switch. Constitutive expression of LacI and CymR, inducible $P_{tac}$*imm2* and $P_{cym}$*colE2.* Only the cytotoxic portion of the *col* gene is included with an added start codon. | This study |
| pTH109 | Tse2-Tsi2 kill switch. Constitutive expression of LacI and CymR, inducible $P_{tac}$*tsi2* and $P_{cym}$*tse2.* | This study |
| pTH121 | EcoRI kill switch with the methylase removed. Constitutive expression of LacI and CymR, inducible expression of $P_{cym}$*ecoRI.* | This study |
| pTH181 | TisB kill switch without an antitoxin. Constitutive expression of LacI and CymR, inducible expression of $P_{cym}$*tisB.* | This study |
| pTH213 | TisB-IstR-1 kill switch. Constitutive expression of LacI and CymR, inducible expression of $P_{tac}$*IstR-2* $P_{cym}$*tisB.* | This study |
| pTH51-GFP-ssrA (LAA) | Derivative of pTH51 kill switch reporter construct with SBW25 native ssrA tag added to the C-terminus of GFP. Produces undetectable levels of GFP fluorescence. | This study |
| pTH51-GFP-ssrA (DAS) | Derivative of pTH51 kill switch reporter construct with SBW25 altered ssrA tag added to the C-terminus of GFP to reduce expression. | This study |

| | | |
|---|---|---|
| pTH51-GFP-ssrA (ILV) | Derivative of pTH51 kill switch reporter construct with SBW25 altered ssrA tag added to the C-terminus of GFP to reduce expression | This study |
| pTH51-GFP-ssrA (LVC) | Derivative of pTH51 kill switch reporter construct with SBW25 altered ssrA tag added to the C-terminus of GFP to reduce expression | This study |
| pTH51-GFP-ssrA (AAV) | Derivative of pTH51 kill switch reporter construct with SBW25 altered ssrA tag added to the C-terminus of GFP to reduce expression | This study |
| pTH193 | Derivative of pTH107. Immunity E2 contains SBW25 ssrA tag ending in -AAV. | This study |
| pTH194 | Derivative of pTH107. Immunity E2 contains SBW25 ssrA tag ending in -LVC. | This study |
| pTH196 | Derivative of pTH107. Immunity E2 contains SBW25 ssrA tag ending in -DAS. | This study |
| pTH197 | Derivative of pTH109. Tsi2 contains SBW25 ssrA tag ending in -AAV. | This study |
| pTH198 | Derivative of pTH107. Tsi2 contains SBW25 ssrA tag ending in -LVC. | This study |
| pTH200 | Derivative of pTH107. Tsi2 contains SBW25 ssrA tag ending in -DAS. | This study |

**Table S3. gBlock sequences of kill switch circuits.** Inactivator genes are highlighted in blue, toxin genes are green. Sequences of modified immunity genes containing a representative *ssrA* tag are also provided, with the *ssrA* sequence highlighted.

| Toxin/Inactivator | Toxin Sequence |
|---|---|
| TisB/IstR | ttgacaattaatcatcggctcgtataatgtgtggaattgtgagcgctcacaattGCACTAAATACGTCAAAATTCGTGCCGAAATTGCGCGTTCTGCGCGGAACACGTATACTTTCAGTGTTGACATAATACAGTGTGCTTTGCGGTTACCAGCCGCAGGCGACTGACGAAACCTCGCTCCGGCGGGGTTTTTTCCCGCACGCTAAATATAAAAGGGGAGCGGTTTCCCGCTCCCCTTTGGTGCGACTTGAATCTGAAttaCTTCAGGTATTTCAGAACAGCATCAAGCAGTTGCAGTGCTGCAACAATGAGTTTGAGGATAAGAATGGCGATATCCACCAGGTTcatACGCGTCTCCTGTGGTTCAGGAGCACTGCCAGCTGACGTACCTTTCCGCTGCCTGTTGCCAGCTGTTGTGCACGCGGTATCTCAAGGAGAAACGTTGTGCTTGTTGACATAACACAGTGTGCTCctctttaattctagaataat |
| CcdB/CcdA | attacgaggcgtatcacgaggccctttcgtcttcacctcgagtgttgacaattaatcatcggctcgtataatgtgtggaattgtgagcgctcacaatttcacacaTCTAGAGCTAATCATCGCGTACTCAGGAGGCAAGTAATGAAGCAGCGGATTACGGTCACCGTCGACAGCGATAGCTATCAGCTCCTGAAAGCGTATGACGTGAATATTTCCGGGCTCGTGTCCACGACCATGCAGAATGAAGCGCGGCGGCTCCGCGCCGAGCGGTGGAAAGCTGAAAATCAGGAGGGCATGGCTGAAGTCGCTCGTTTTATTGAAATGAACGGTTCCTTTGCAGACGAGAACCGCGACTGGTAGgaggatccagatctcatcaccatcaccatcactaagcttaattagctgagcttggactcctgttgatagatccagtaatgacctcagaactccatctggatttgttcagaacgctcggttgccgccgggcgttttttattggtgagaatccaagctagcttggcgagatggagtccgaggcctcgagatgcttaaagcggccgcCTAGATACCCCAGAACATCAAATTGATTGCGTTTTTAATGTCGTTCTCCCGATGGCTCAGGTCCGCGACTTCTTCCCCAATCACGCTCACGGGGACGGATGCCATATCGGTGGTCATCATGCGCCAGCTCTCGTCGCCAATGTGGACCACTGGATACAGCTCGCGGGACACTTTATCGCTCAACAACCGCGCGGACGCGAGGGGAATGACCATACGGCGACCCGGGGTGTCAATGATATCGCTCTGGACATCCACGAACAGCCGGTACCGGGATTCACGCTTATAGGTGTACACCTTGAATTGCATggttaatttctcctctttaattctagaataat |
| HicA/HicB | attacgaggcgtatcacgaggccctttcgtcttcacctcgagtgttgacaattaatcatcggctcgtataatgtgtggaattgtgagcgctcacaatttcacacaTCTAGAGCTAATCATCGCGTACTCAGGAGGCAAGTAATGCGTTATCCCGTCACTCTTACACCCGCGCCGGAAGGCGGTTATATGGTTTCTTTTGTGGATATCCCTGAAGCGTTGACCCAGGGCGAAACTGTCGCTGAAGCGATGGAAGCGGCAAAAGATGCTTTACTGACCGCATTTGATTTTTATTTTGAAGATAACGAGCTTATCCCTTTACCTTCGCCATTAAATAGTCACGATCACTTTATTGAAGTACCTTTGAGCGTCGCCTCTAAGGTATTGCTGTTAAATGCTTTTTTACAGTCAGAAATCACTCAGCAAGAGTTAGCCAGGCGAATTGGCAAACCTAAACAGGAGATTACTCGCCTATTTAACTTGCATCATGCGACAAAAATCGACGCCGTCCAGCTCGCGGCAAAGGCGCTTGGCAAAGAGTTATCGCTGGTGATGGTTTAAgaggatccagatctcatcaccatcaccatcactaagcttaattagctgagcttggactcctgttgatagatccagtaatgacctcagaactccatctggatttgttcagaacgctcggttgccgccgggcgttttttattggtgagaatccaagctagcttggcgagatggagtccgaggcctcgagatgcttaaagcggccgcTTAACTCAAACCGAGTTGTTTCAGGATTGCTTTACGCAATGGTTCTTTAATCTCATCGCAGGGGTGACGCGGCATGACACTGCGCCTCCCATGAAACCTGAGTTTCAAATGGTTGCTGCCATTCGCTACATCGACGCCCTGAGATTCGAGCCAACGTCTGAACTCGCTTTGTTTCACggttaatttctcctctttaattctagaataat |

| | |
|---|---|
| ParE/ParD | attacgaggcgtatcacgaggccctttcgtcttcacctcgagtgttgacaattaatcatcggctcgtataatgtgtgga<br>attgtgagcgctcacaatttcacacaTCTAGAGCTAATCATCGCGTACTCAGGAGGCAAGT<br>AATGTCCCGCCTCACCATTGACATGACCGATCAGCAGCACCAAAGCCTGAAG<br>GCGTTGGCGGCATTGCAGGGCAAAACCATTAAGCAATACGCCCTGGAACGTT<br>TGTTCCCCGGTGATGCCGACGCTGACCAGGCATGGCAGGAGTTGAAGACCA<br>TGCTCGGTAACCGGATCAACGACGGTCTGGCCGGTAAAGTCAGCACCAAATC<br>GGTCGGGGAAATCTTGGACGAAGAGCTGTCGGGGGATCGTGCGTAAgaggatc<br>cagatctcatcaccatcaccatcactaagcttaattagctgagcttggactcctgttgatagatccagtaatgacctca<br>gaactccatctggatttgttcagaacgctcggttgccgccgggcgttttttattggtgagaatccaagctagcttggcg<br>agatggagtccgaggcctcgagatgcttaaagcggccgcTTAGCCCTTGAGCCGATCAGCGAG<br>ACGCGTCATCAGATCCATCCGTTCATGCAAAATAGCCACCACGAGTGCCGGT<br>TCGCCTGCGCGAGGCAAGCAGAAGACATAGTGATGCTCACACCGTGCCATG<br>CGCAAAGCGGGGAAGAGTTCGGACATATCCTTGAAAGGCCCTTCGCCGGCT<br>GCCAGACGCGCAATCCCCTGCTCCAATTTCGCAATATAGCGGCGGACCTGCG<br>CTGCCCCCCACTCACGGCGGGTGTAACGGATAATGCCACGGAGATCAGCTT<br>CAGCCTCTGCGGTCAGAATATATGCGGTCATggttaatttctcctctttaattctagaataat |
| RelE/RelB | attacgaggcgtatcacgaggccctttcgtcttcacctcgagtgttgacaattaatcatcggctcgtataatgtgtgga<br>attgtgagcgctcacaatttcacacaTCTAGAGCTAATCATCGCGTACTCAGGAGGCAAGT<br>AATGGGTTCGATCAACCTGCGGATTGACGACGAACTGAAAGCCCGGAGCTAC<br>GCAGCCCTGGAGAAGATGGGGGTGACGCCATCCGAAGCCCTGCGCTTGATG<br>TTGGAATACATTGCTGACAATGAACGGTTGCCCTTCAAACAGACGCTGCTGTC<br>CGATGAGGACGCCGAGTTGGTCGAGATCGTGAAGGAACGGTTGCGGAATCC<br>AAAGCCTGTGCGGGTGACCTTGGACGAGTTGTAGgaggatccagatctcatcaccatcac<br>catcactaagcttaattagctgagcttggactcctgttgatagatccagtaatgacctcagaactccatctggatttgtt<br>cagaacgctcggttgccgccgggcgttttttattggtgagaatccaagctagcttggcgagatggagtccgaggcct<br>cgagatgcttaaagcggccgcCTACAGGATACGCTTCACCGCCTCCGAGTAGACTTCC<br>GACCGCTCCCGCTTCCCCACGGAGATGACGAACACCACCACCTTCTCATCAA<br>TCACCTGGTAGACCAGCCGATAACCCGAGGAACGGAGTTTGATTTTGTAACA<br>ATCCGGCATGCCACGGAGCTTATTTGCTTCAATGCGAGGGCTTTCCAGCACT<br>TCGACGAGCTTTTTTTTCAACTGCTCGCGCACGGTCGAACCGAGCTTACGCC<br>ATTCTTTCAGTGCCCGTTCGTCAAAGTCCAGAAAATAGGCCATggttaatttctcctcttt<br>aattctagaataat |
| Colicin E2/<br>Immunity E2 | attacgaggcgtatcacgaggccctttcgtcttcacctcgagtgttgacaattaatcatcggctcgtataatgtgtgga<br>attgtgagcgctcacaatttcacacaTCTAGAGCTAATCATCGCGTACTCAGGAGGCAAGT<br>Aatggaactgaaacatagtattagtgattataccgaggctgaatttctggagtttgtaaaaaaaatatgtagagctga<br>aggtgctactgaagaggatgacaataaattagtgagagagtttgagcgattaactgagcacccagatggttcagat<br>ctgatttattatcctcgcgatgacagggaagatagtcctgaagggattgtcaaggaaattaaagaatggcgagctg<br>ctaacggtaagtcaggatttaaacagggctgagaggatccagatctcatcaccatcaccatcactaagcttaatta<br>gctgagcttggactcctgttgatagatccagtaatgacctcagaactccatctggatttgttcagaacgctcggttgcc<br>gccgggcgttttttattggtgagaatccaagctagcttggcgagatggagtccgaggcctcgagatgcttaaagcgg<br>ccgcttacttaccccgatgaatatcaatatgtcgcttaggtgtggtcactctgatattattcatatcatagacaccaccat<br>cctgactgattggtttatcatgatgtaattcaaagcgttccctaccacctacttggtctttcttccttgcaaaaggtgcttttc<br>ccttttgaatgttcgtcttattactgcctttaaattgcttactaagatcgggatcttttgacacttcttcccagaatttcttccgg<br>aaatcgtcaaagtttttaaattctttatcacgcaacttatcagcaatgcgatctggaattggcgctcctgaatctttacct<br>gcatcatccagccatttatcaccaactggtttacctttacctgtcgccttcccCATtaaggttaatttctcctctttaattct<br>agaataat |
| Tse2/Tsi2 | Attacgaggcgtatcacgaggccctttcgtcttcacctcgagtgttgacaattaatcatcggctcgtataatgtgtgga<br>attgtgagcgctcacaatttcacacaTCTAGAGCTAATCATCGCGTACTCAGGAGGCAAGT<br>AATGAACCTGAAACCCCAGACCCTGATGGTGGCGATCCAGTGCGTCGCCGC<br>GCGCACCCGCGAACTCGACGCGCAGTTGCAGAACGACGACCCGCAGAACGC<br>CGCTGAACTCGAACAGTTGCTGGTCGGCTACGACCTCGCCGCCGACGACCT<br>GAAGAACGCCTACGAGCAAGCCCTGGGCCAATACAGCGGCCTGCCGCCCTA<br>TGACCGGCTGATCGAAGAGCCGGCATCCTGAgaggatccagatctcatcaccatcaccatc |

| | |
|---|---|
| | actaagcttaattagctgagcttggactcctgttgatagatccagtaatgacctcagaactccatctggatttgttcaga<br>acgctcggttgccgccgggcgtttttattggtgagaatccaagctagcttggcgagatggagtccgaggcctcgag<br>atgcttaaagcggccgcCTACAGGCCCCGTGCTTTCTCCCACTGGCGGCGAATAGCG<br>TTACGGACGAAGTTGTCCAGCTGGCCCATCAAGACTTCCCGATACATAGGTTT<br>AGCGGGCATAATCGTGTAGTGGGTAGCTTGCAGACGTTTATTATAGCTATCCT<br>GCTTCACCTTCAAGTCAGGAGGAATGTCCGTCCCATCCGGAATGACAAAGTC<br>ACCGTCAGCCCGCCGCAACACACCGGCACGATCGAAGACGGACGTACCACC<br>CGTATCTGGGGCATAGAGACCATCCCGACCTGCGACGATATCTGGGGCACG<br>AGCAGGCACGCCAGGGGTATTTGGGCTATCAAAGGTCGGATGCAACAGCGG<br>CGCCACTTCTGCCGCTTCGGCGAGTTCTTTGTCCGGGTGCAGATAACGCCCG<br>ACGTCCCCATCGTAATTTGCTTTGAAGACAGCACGATACAACGTGAGCGACG<br>TCTTTTCGTAGTCATAGCTCATggttaatttctcctctttaattctagaataat |
| EcoRI/EcoRIM | attacgaggcgtatcacgaggccctttcgtcttcacctcgagtgttgacaattaatcatcggctcgtataatgtgtgga<br>attgtgagcgctcacaatttcacacaTCTAGAGCTAATCATCGCGTACTCAGGAGGCAAGT<br>AATGGCACGTAATGCAACGAACAAACTGTTGCACAAGGCTAAGAAATCGAAAA<br>GCGATGAATTTTATACGCAGTATTGCGATATCGAAAATGAATTGCAGTACTAC<br>CGTGAACATTTCAGCGACAAGGTGGTGTATTGTAATTGTGATGATCCGCGTGT<br>CTCCAACTTCTTTAAGTATTTCGCCGTCAACTTCGATAACCTGGGGTTGAAGA<br>AACTGATCGCGAGCTGCTATGTCGAAAACAAGGAAGGTTTCTCCTCCTCCGA<br>GGCTGCGAAGAATGGCTTTTACTACGAGTACCACAAGGAGAATGGCAAAAAA<br>CTCGTGTTCGACGATATCTCCGTCAGCAGCTTCTGCGGGGACGGGGATTTTC<br>GCAGCAGCGAGAGCATTGACCTCCTGAAGAAATCGGATATCGTCGTGACCAA<br>TCCACCATTTTCCCTCTTCCGCGAATATCTCGACCAGCTCATCAAATATGATAA<br>AAAATTTCTCATTATTGCAAATGTGAACAGCATCACGTATAAGGAGGTGTTCAA<br>TCTCATCAAGGAAAACAAGATTTGGCTCGGGGTGCACCTCGGCCGTGGGGTC<br>TCGGGGTTTATTGTGCCTGAACACTATGAGCTCTACGGGACGGAAGCCCGGA<br>TTGATTCGAATGGGAATCGCATCATTAGCCCAAACAACTGCCTGTGGCTCAC<br>GAATCTCGACGTGTTCATTCGTCACAAAGATTTGCCGCTCACGCGGAAATACT<br>TTGGCAATGAAAGCTCGTACCCAAAATACGACAACTATGATGCCATCAATGTC<br>AACAAGACGAAGGATATCCCACTGGACTATAATGGCGTCATGGGTGTGCCGA<br>TTACGTTCTTGCACAAGTTTAACCCCGAACAATTTGAGTTGATTAAATTTCGTA<br>AAGGTGTCGACGAAAAAGACCTGTCGATCAATGGTAAATGTCCATATTTCCGC<br>ATTTTGATCAAGAATAAACGTTTGCAGAAGTAGgaggatccagatctcatcaccatcaccat<br>cactaagcttaattagctgagcttggactcctgttgatagatccagtaatgacctcagaactccatctggatttgttcag<br>aacgctcggttgccgccgggcgtttttattggtgagaatccaagctagcttggcgagatggagtccgaggcctcga<br>gatgcttaaagcggccgcCTATTTGGAGGTGAGCTGTTCGAAGAGATCCCGCCCCAG<br>CACCCGCAGCGACGTCGTGGAGATATCGAACATGATTTCGAACATGATCTTG<br>GAATCCCATTCCCGACCGTCCCCTTGGGTATAAATCGAAGCTGCTTGCAACAT<br>GATCGACTTGTCCTTATGATTGACGAACTTGTTAATACACAAGTTCGAATTAAT<br>TGGCATACCGTAATTTGCAGCCGTCAGGCGGTCCAAACGGTTCAAGATACCC<br>GAATTATACTCGAGGTTCACGACGCGACCATCAGGCCGGGTAATCGAGATGT<br>TTTCGGTGAGGAAATTCGAACCCTCCAGAAACAGCACATATGGAAAGTGGCT<br>CTCGGAGAGCATAAAATTCGCAATCTCCGAAATATTTTTGTGGGACCGCTCGA<br>TTGCATTCCCCGCCGCCATCAAGTCTTGGTCCCCCCGCTTCCCCACGAGCAG<br>GCCATTACGGATATTAATAATGTCCTTACCTTGGTGTTTCGCCTCTGCGACCA<br>AGACGACACGCCATTCGCCATAATCATCTTTGACTTCGACGATCCCACCGTCA<br>GGTTTAATGCTCGAATTGGAGACGAACAACGTACCGCCGAGATCAGGGTCAA<br>TTTTTTTCAGAGCCTCATTGATCTCGGTCTTTTTAATCGAGTCGCGATACCGAA<br>AGGAGAGCTGTGGGTACTCATTCGACAAAGCCTTCTTCACCAGCTTCGACAC<br>TTCCCCGACCGCCAAGTCATGGGCCTTAGCATAGTCACCGAAAATCCCAATG<br>ACACCCTGGGACAACTTGTGTTGCTCGGTCAGACGGTTCGACTGTTTTTTATT<br>GCTCATggttaatttctcctctttaattctagaataat |
| TisB | gactcctgttgatagatccagtaatgacctcagaactccatctggatttgttcagaacgctcggttgccgccgggcgtt<br>ttttattggtgagaatccaagctagcttggcgagatggagtccgaggcctcgagatgcttaaagcggccgcTTAC |

| | |
|---|---|
| | TTCAAGTACTTGAGCACGGCATCCAACAGTTGCAATGCAGCGACAATCAGCTTGAGGATCAAAATAGCGATATCGACCAGGTTCATggttaatttctcctctttaattctagaataat |
| Immunity E2-SsrA[AAV] | attctcaccaataaaaaaacgcccggcggcaaccgagcgttctgaacaaatccagatggagttctgaggtcattactggatctatcaacaggagtccaagctcagctaattaagcttagtgatggtgatggtgatgagatctggatcctcctaCACTGCAGCGGCGAACTCCTGGCCGTAGTTCTCGTCGTTGGCgccctgtttaaatcctgacttaccgttagcagctcgccattctttaatttccttgacaatcccttcaggactatcttccctgtcatcgcgaggataataaatcagatctgaaccatctgggtgctcagttaatcgctcaaactctctcactaatttattgtcatcctcttcagtagcaccttcagctctacatattttttttttacaaactccagaaattcagcctcggtataatcactaatactatgtttcagttccatTACTTGCCTCCTGAGTACGCGATGATTAGCTCTAGAtgtgtgaaattgtgagcgctcacaattccacacattatacgagccgatgattaattgtcaa |
| Tsi2-SsrA[AAV] | attctcaccaataaaaaaacgcccggcggcaaccgagcgttctgaacaaatccagatggagttctgaggtcattactggatctatcaacaggagtccaagctcagctaattaagcttagtgatggtgatggtgatgagatctggatcctcctaCACCGCAGCGGCGAACTCCTGGCCGTAGTTCTCGTCGTTGGCggatgccggctcttcgatcagccggtcatagggcggcaggccgctgtattggcccagggcttgctcgtaggcgttcttcaggtcgtcggcggcgaggtcgtagccgaccagcaactgttcgagttcagcggcgttctgcgggtcgtcgttctgcaactgcgcgtcgagttcgcgggtgcgcgcggcgacgcactggatcgccaccatcagggtctggggtttcaggttcatTACTTGCCTCCTGAGTACGCGATGATTAGCTCTAGAtgtgtgaaattgtgagcgctcacaattccacacattatacgagccgatgattaattgtcaa |

#### Table S4. Escape mutations for all circuits.

| | Locus | Mode | Consequence | Mechanism |
|---|---|---|---|---|
| CcdB-CcdA | | | | |
| 1 | LacI | Duplication | Frameshift | TGGC duplication at +605 (TCTGGCTGGCTGGCATAAATATCT > TCTGGCTGGCTGGCTGGCATAAATATCT) |
| 2 | LacI | Duplication | Frameshift | TGGC duplication at +605 (TCTGGCTGGCTGGCATAAATATCT > TCTGGCTGGCTGGCTGGCATAAATATCT) |
| 3 | LacI | Duplication | Frameshift | TGGC duplication at +605 (TCTGGCTGGCTGGCATAAATATCT > TCTGGCTGGCTGGCTGGCATAAATATCT) |
| 4 | LacI | Deletion | Frameshift | TGGC deletion at +601, 212STOP (TCTGGCTGGCTGGCATAAATATCT > TCTGGCTGGC----ATAAATATCT) |
| 4 | Ori | Indel | - | GCCTCCCCCGCCCC > GCCTCCCCCCGCCCC |
| 5 | LacI | Substitution | Stop codon | E259 > STOP |
| 6 | LacI | Duplication | Frameshift | TGGC duplication at +605 (TCTGGCTGGCTGGCATAAATATCT > TCTGGCTGGCTGGCTGGCATAAATATCT) |
| 7 | Toxin | Deletion | Stop codon | 13 bp deletion at R86 leading to C-terminal frameshift beginning at I90 |
| HicA-HicB | | | | |
| 1 | LacI | Duplication | Frameshift | TGGC duplication at +605 (TCTGGCTGGCTGGCATAAATATCT > TCTGGCTGGCTGGCTGGCATAAATATCT) |
| 2 | Toxin | Indel | Frameshift | L52 > STOP |
| 3 | LacI | Duplication | Frameshift | 25bp duplication at Leu233, 243 > STOP |
| 4 | LacI | Duplication | Frameshift | TGGC duplication at +605 (TCTGGCTGGCTGGCATAAATATCT > TCTGGCTGGCTGGCTGGCATAAATATCT) |
| 5 | LacI | Duplication | Frameshift | TGGC duplication at +605 (TCTGGCTGGCTGGCATAAATATCT > TCTGGCTGGCTGGCTGGCATAAATATCT) |
| 6 | LacI | Duplication | Frameshift | TGGC duplication at +605 (TCTGGCTGGCTGGCATAAATATCT > TCTGGCTGGCTGGCTGGCATAAATATCT) |
| 7 | Toxin | Deletion | Deletion | 147 bp deletion including most of HicA + half of RBS |
| ParE-ParD | | | | |
| 1 | Toxin | Indel | Frameshift | Duplication of G (+79) |
| 2 | Entire circuit | Deletion | No expression | Deletion of most of insert (2484 bp total, starts at M54 in ParE, ParD, CymR, and N-terminal portion of LacI.) |
| 3 | Toxin | Deletion | Premature stop | 175bp deletion starts at Ala85 and stretches into terminator, resulting in truncated protein. |
| 4 | Toxin | Indel | Frameshift | Duplication of G (+79) |
| 5 | Toxin | Substitution | Nonsense mutation | C > T transition leading to Q39 > STOP. |
| 6 | Toxin | Deletion | Frameshift | 159bp deletion +130 into ParE. |
| 7 | Toxin | Deletion | Premature stop | 49bp deletion +11 into ParE. |

Table S4. cont'd

| | Locus | Mode | Consequence | Mechanism |
|---|---|---|---|---|
| Tse2-Tsi2 | | | | |
| 1 | LacI | Substitution | Stop codon | W201 > STOP |
| 2 | LacI | Substitution | Stop codon | W201 > STOP |
| 2 | Pcym | Deletion | Removal of promoter | Operator sites deleted by homology (-43:-101) |
| 3 | Pcym | Deletion | Removal of promoter | Operator sites deleted by homology (-43:-101) |
| 3 | Ori | Indel | - | GCCTCCCCGCCCC > GCCTCCCCCGCCCC |
| 4 | Pcym | Deletion | Removal of promoter | Operator sites deleted by homology (-43:-101) |
| 5 | Pcym | Deletion | Removal of promoter | Operator sites deleted by homology (-43:-101) |
| 6 | Pcym | Deletion | Removal of promoter | Operator sites deleted by homology (-43:-101) |
| 6 | Ori | Indel | - | GCCTCCCCGCCCC > GCCTCCCCCGCCCC |
| 7 | Pcym | Substitution | Change in -10 sequence | A > G transition |
| Col-ImmE2 | | | | |
| 1 | LacI | Duplication | Frameshift | TGGC duplication at +605 (TCTGGCTGGCTGGCATAAATATCT > TCTGGCTGGCTGGCTGGCATAAATATCT) |
| 2 | LacI | Duplication | Frameshift | TGGC duplication at +605 (TCTGGCTGGCTGGCATAAATATCT > TCTGGCTGGCTGGCTGGCATAAATATCT) |
| 2 | Ori | Indel | - | GCGTTTTTTGCGTTTCCA > GCGTTTTTTTGCGTTTCCA |
| 3 | LacO | Substitution | - | G > A transition in operator (-53) |
| 4 | Pcym | Substitution | - | A > G transition in promoter (-71) |
| 5 | LacO | Substitution | - | C > T transition in operator (-54) |
| 6 | LacI | Duplication | Frameshift | TGGC duplication at +605 (TCTGGCTGGCTGGCATAAATATCT > TCTGGCTGGCTGGCTGGCATAAATATCT) |
| 7 | LacO | Substitution | - | G > C transversion in operator (-59) |
| RelE-RelB | | | | |
| 1 | Toxin | Deletion | No RelE expression | Deletion of -112 (Pcym promoter) into RelE |
| 2 | Toxin | Substitution | Premature stop | G > transition resulting in Q65 > STOP |
| 3 | LacI | Deletion | Frameshift | Deletion of 5nt after G121 (+363) resulting in truncated protein of 121 aa. |
| 4 | LacI | Indel | Frameshift | Insertion of C after I268 (+804) leading to truncated protein of 278aa (wild type is 360). |
| 6 | LacI | Duplication | Frameshift | Duplication of CCAG after R197 (+591) leading to truncated protein of 203 aa. |
| 7 | LacI | Duplication | Frameshift | Duplication of bp 49-56 resulting in insertion of 3 AA's (YMI) after S15 and a truncated protein of 85 aa. |

Table S4. cont'd

| | Locus | Mode | Consequence | Mechanism |
|---|---|---|---|---|
| Col-ImmE2(ssrA) | | | | |
| 1 | LacI | Duplication | Frameshift | TGGC duplication at +605 (TCTGGCTGGCTGGCATAAATATCT > TCTGGCTGGCTGGCTGGCATAAATATCT) |
| 2 | LacI | Duplication | Frameshift | TGGC duplication at +605 (TCTGGCTGGCTGGCATAAATATCT > TCTGGCTGGCTGGCTGGCATAAATATCT) |
| 3 | LacI | Duplication | Frameshift | TGGC duplication at +605 (TCTGGCTGGCTGGCATAAATATCT > TCTGGCTGGCTGGCTGGCATAAATATCT) |
| 4 | LacI | Duplication | Frameshift | TGGC duplication at +605 (TCTGGCTGGCTGGCATAAATATCT > TCTGGCTGGCTGGCTGGCATAAATATCT) |
| 5 | LacI | Duplication | Frameshift | TGGC duplication at +605 (TCTGGCTGGCTGGCATAAATATCT > TCTGGCTGGCTGGCTGGCATAAATATCT) |
| 7 | LacI | Duplication | Frameshift | TGGC duplication at +605 (TCTGGCTGGCTGGCATAAATATCT > TCTGGCTGGCTGGCTGGCATAAATATCT) |
| Tse2-Tsi2(ssrA) | | | | |
| 1 | LacI | Duplication | Frameshift | TGGC duplication at +605 (TCTGGCTGGCTGGCATAAATATCT > TCTGGCTGGCTGGCTGGCATAAATATCT) |
| 2 | LacI | Duplication | Frameshift | TGGC duplication at +605 (TCTGGCTGGCTGGCATAAATATCT > TCTGGCTGGCTGGCTGGCATAAATATCT) |
| 3 | LacI | Duplication | Frameshift | TGGC duplication at +605 (TCTGGCTGGCTGGCATAAATATCT > TCTGGCTGGCTGGCTGGCATAAATATCT) |
| 4 | LacI | Duplication | Frameshift | TGGC duplication at +605 (TCTGGCTGGCTGGCATAAATATCT > TCTGGCTGGCTGGCTGGCATAAATATCT) |
| 5 | LacI | Indel | Frameshift | Deletion of +C81, resulting in truncated peptide. |
| 6 | LacI | Duplication | Frameshift | TGGC duplication at +605 (TCTGGCTGGCTGGCATAAATATCT > TCTGGCTGGCTGGCTGGCATAAATATCT) |
| 7 | LacI | Substitution | Missense mutation | G252S |

Table S4. cont'd

| | Locus | Mode | Consequence | Mechanism |
|---|---|---|---|---|
| EcoRI (no methylase) | | | | |
| 1 | Toxin | Deletion | In-frame deletion | In-frame 12bp deletion (A758-A770) at +747 resulting in E253D and removeal of I254-D257 |
| 2 | Toxin | Substitution | Missense mutation | D133G |
| 3 | Toxin | Duplication | In-frame duplication | 12bp duplication (AATCATGTTCGA) at D257 resulting in D257E and insertion of IMFD (258-261). |
| 4 | Toxin | Duplication | In-frame duplication | 12bp duplication (AATCATGTTCGA) at D257 resulting in D257E and insertion of IMFD (258-261). |
| 5 | Toxin | Deletion | Frameshift | 100bp deletion (+171 - 271), 103 > STOP |
| 6 | Toxin | Deletion | Frameshift | 10 bp deletion (+561 - 571), 190 > STOP |
| 7 | Toxin | Substitution | Nonsense mutation | Q240 > STOP |
| TisB (no antitoxin) | | | | |
| 1 | Pcym | Deletion | - | Operator sites deleted by homology (-43:-101) |
| 2 | Pcym | Deletion | - | Operator sites deleted by homology (-43:-101) |
| 3 | Pcym | Deletion | - | Operator sites deleted by homology (-43:-101) |
| 4 | Pcym | Substitution | - | T > C transition in -10 sequence (-69) |
| 5 | Pcym | Deletion | - | Operator sites deleted by homology (-43:-101) |
| 6 | Pcym | Substitution | - | T > C transition in -10 sequence (-69) |
| 7 | Pcym | Deletion | - | Operator sites deleted by homology (-43:-101) |
